# Genomics of cold adaptations in the Antarctic notothenioid fish radiation

**DOI:** 10.1101/2022.06.08.494096

**Authors:** Iliana Bista, Jonathan M. D. Wood, Thomas Desvignes, Shane A. McCarthy, Michael Matschiner, Zemin Ning, Alan Tracey, James Torrance, Ying Sims, William Chow, Michelle Smith, Karen Oliver, Leanne Haggerty, Walter Salzburger, John H. Postlethwait, Kerstin Howe, Melody S. Clark, William H. Detrich, C.-H. Christina Cheng, Eric A. Miska, Richard Durbin

## Abstract

Numerous novel adaptations characterise the radiation of notothenioids, the dominant fish group in the freezing seas of the Southern Ocean. To improve understanding of the evolution of this iconic fish group, we generated and analysed new genome assemblies for 24 species covering all major subgroups of the radiation. We present a new estimate for the onset of the radiation at 10.7 million years ago, based on a time-calibrated phylogeny derived from genome-wide sequence data. We identify a two-fold variation in genome size, driven by expansion of multiple transposable element families, and use long-read sequencing data to reconstruct two evolutionarily important, highly repetitive gene family loci. First, we present the most complete reconstruction to date of the antifreeze glycoprotein gene family, whose emergence enabled survival in sub-zero temperatures, showing the expansion of the antifreeze gene locus from the ancestral to the derived state. Second, we trace the loss of haemoglobin genes in icefishes, the only vertebrates lacking functional haemoglobins, through complete reconstruction of the two haemoglobin gene clusters across notothenioid families. Finally, we show that both the haemoglobin and antifreeze genomic loci are characterised by multiple transposon expansions that may have driven the evolutionary history of these genes.

## Introduction

The suborder Notothenioidei is a textbook example of a marine fish adaptive radiation, with notothenioids being the dominant fish group of the Southern Ocean, both in terms of species richness and biomass, comprising between 130-140 species^1–3^ (**Fig. 1A**). The establishment and initial diversification of the notothenioids is closely linked to the separation of the Antarctic continent from surrounding land masses and the subsequent establishment of the Antarctic Circumpolar Current (ACC)^4^ (**Fig. 1B**). Formation of the ACC exacerbated the isolation of the Antarctic continent and contributed to cooling of the surrounding waters, glaciation of the continent, and appearance of sea ice^5^. These events in turn extirpated most of the original temperate fish fauna, and notothenioids expanded to fill the abandoned niches as they evolved adaptations to life in the isolated, cold, and highly oxygenated waters of the Southern Ocean^6,7^. Since notothenioids include species occurring in cool-temperate non-Antarctic regions^8^, as well as species occurring in icy, freezing higher latitudes (known as the “Antarctic clade” or Cryonotothenioidea (cryonotothenioids))^9^, they represent a powerful model for the study of the genomic origins of extremophiles. Adaptations to cold include the presence of antifreeze glycoproteins (AFGPs)^10^, the lack of the classic heat shock response^11^, and the presence of giant muscle fibres in some notothenioids^12^. Also, a deleterious, but non-lethal, respiratory phenotype arose in the derived family Channichthyidae (“icefishes”), including the complete loss of functional haemoglobins in all of its species and the loss of cardiac myoglobin in six of them^13^. While haemoglobin loss was not lethal in the oxygenated waters of the cold Southern Ocean, these losses were not without fitness consequences, as illustrated by numerous compensatory cardiovascular adaptations, including enlarged hearts, and increased vascular bores^49^.

**Figure 1.**
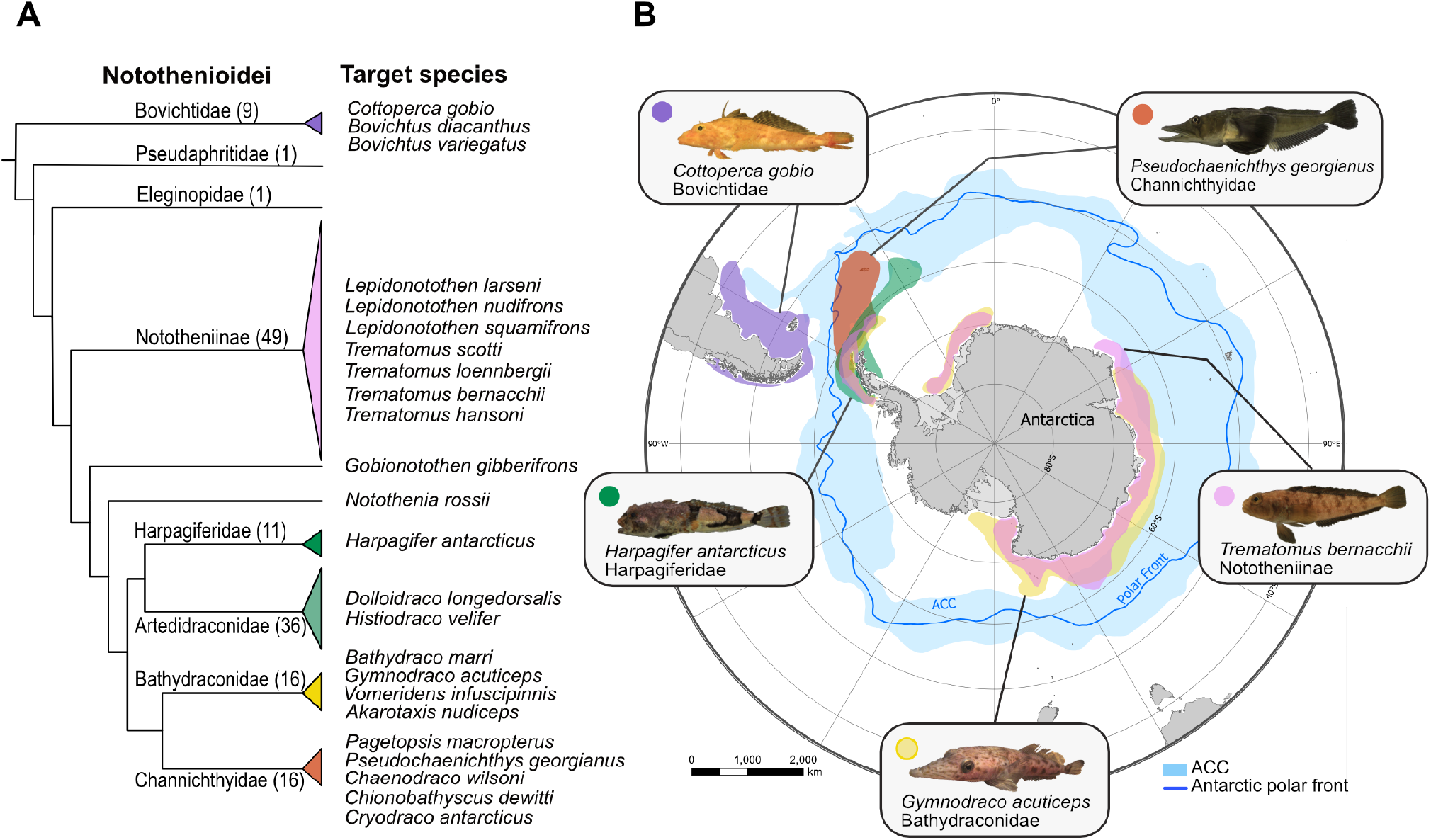
The Notothenioidei suborder, all 24 target species, and representative geographic distributions of PacBio sequenced species. A) Notothenioid families and number of species contained in each family based on the species list by ref^3^. The target species sequenced in the present study are listed, and the number of species per family is shown in parenthesis, except for the Nototheniidae which are paraphyletic, containing the Nototheniinae, *G. gibberifrons* and *N. rossi.* B) Map of Antarctica and the Southern Ocean showing the geographic distribution of the five notothenioid species sequenced with PacBio. Colours correspond to the respective families on the tree. The blue belt around Antarctica indicates the region of the Antarctic Circumpolar Current (ACC), and the thin blue line the Antarctic polar front. Data for geographic distribution of each species were extracted from the SCAR Antarctic Biodiversity Portal, comprising fishing records from multiple databases.

The in-depth characterisation of notothenioid genomes has been hampered in the past by their complex genome characteristics, such as high levels of repeats and heterozygosity, that have hindered the accuracy of genome assemblies based on short-read data. Furthermore, the few available high quality notothenioid genome assemblies^14,15^ cover only a small portion of the diversity in this group. The Vertebrate Genomes Project (VGP) (https://vertebrategenomesproject.org/)^16^ has demonstrated that long-read sequencing technologies can generate highly contiguous genome assemblies even for the most technically difficult species. To produce a step-change increase in Antarctic notothenioid genome resources for the broader community, we have applied the VGP pipeline and standards to five selected notothenioid species representing key points in the radiation, extended with other sequencing approaches to a total of 24 new genomes. Collectively they cover six of the eight notothenioid families (all except two single-species families) (**Fig. 1A**), including the five families that comprise the Antarctic radiation, and a non-Antarctic family. Using these new genome assemblies, we address previously intractable questions about the evolution of *afgp* genes, the loss of haemoglobins in icefish, and the role of transposable elements in cold adaptation throughout this adaptive radiation.

### Genome sequencing, assembly, and annotation

We generated and analysed reference genome assemblies for 24 notothenioid fish species across the radiation using a variety of sequencing technologies (**Fig. 1, Extended Data Table 1**). Genomes of five species - *Cottoperca gobio* (synonymised by many to *Cottoperca trigloides*^3^), *Trematomus bernacchii, Harpagifer antarcticus, Gymnodraco acuticeps, Pseudochaenichthys georgianus* - each representing a different notothenioid family, were assembled using Pacific Biosciences (PacBio) long reads, in combination with 10X Genomics linked reads and Hi-C data. Briefly, we used Falcon-unzip^17^ to generate each primary assembly from the PacBio reads and then applied 10X Genomics Chromium data for scaffolding and polishing. The genomes of *C. gobio, H. antarcticus* and *P. georgianus* were further scaffolded using Bionano hybrid scaffolding. Additionally for *C. gobio* and *P. georgianus* we also used Hi-C data (ARIMA and Dovetail respectively) for scaffolding with SALSA2^18^. To further improve the quality of these genomes, we performed manual curation using the Genome Evaluation Browser (gEVAL)^19^ to remove mis-assemblies, false duplications, and sequencing contaminations such as symbionts and adapter sequences, and to merge scaffolds based on supporting evidence^20^ (**Supplementary Methods**, **Supplementary Table ST2**). The genomes of *C. gobio* and *P. georgianus*, were assigned to 24 chromosomes, consistent with their known karyotypes^21,22^ (**Extended Data Fig. 1A-B**). The reference assembly for *C. gobio* (fCotGob3.1, GCA_900634415.1) was previously described in ref^23^. Genomes of 11 more species were sequenced with 10X Genomics using a single linked-read library for each and assembled with Supernova v2.0^24^. Genomes of eight additional species were sequenced using only Illumina HiSeq reads and assembled using a reference-guided approach. For these eight species, a primary assembly was generated with SOAPdenovo2^25^, and scaffolding was done using the closest PacBio genome as reference (**Methods, Supplementary Table ST3**). All 19 assemblies were also manually curated to remove external contamination, and false duplications (the latter in Supernova assemblies).

For the PacBio assemblies, BUSCO^26^ gene completeness averages 95% (**Extended Data Fig. 1C**). Gene prediction was performed by Ensembl for all PacBio assemblies via the Ensembl Gene Annotation system^27^ (**Methods, Supplementary Methods**). The BUSCO completeness of the gene annotation averages 92% (**Extended Data Fig. 1D**). Approximately 23-24,000 genes were annotated for the four cryonotothenioid species (24,373 for *T. bernacchii*, 23,146 for *H. antarcticus*, 24,091 for *G. acuticeps*, and 23,222 for *P. georgianus* (**Supplementary Table ST4**)), around 2,000 genes more than the non-Antarctic species *C. gobio* (Ensembl genes: 21,662^23^), suggesting that cold adaptation was accompanied by an expansion in the number of genes in notothenioids (**Extended Data Fig. 1E-F)**.

### A new time-calibrated phylogeny for notothenioids

Our multi-species dataset affords the opportunity to establish a new time-calibrated phylogeny for the notothenioid radiation based on genome-wide data, to help resolve controversies about the timing of the diversification of the notothenioids relative to the chilling of the Southern Ocean. Most of the previously published phylogenetic hypotheses for notothenioids were based on limited numbers of genes^6^, RAD-seq^28,29^, or exome capture data^30^. By analysing genome-wide data extracted from BUSCO single copy genes from 41 percomorph fish species, including the 24 new and eight previously published notothenioid genomes^31,32^ we provide the most comprehensive phylogenomic analyses of notothenioids to date, with taxonomic coverage of most of their sub-groups. Based on this analysis, calibrated using established teleost divergence dates^33^, the onset of diversification of the Cryonotothenioidea, which are characterised by the presence of AFGPs, is estimated at around 10.7 million years ago (MYA) (highest posterior density interval: 14.1–7.8 MYA) (**Fig. 2A**). While the appearance of AFGPs has previously been estimated at 42–22 MYA, which would predate the major cooling of the Southern Ocean^6^, our new analysis indicates that the emergence of AFGPs occurred between 26.3–10.7 MYA. This period includes the Middle Miocene Climate Transition at 15-13 MYA and the subsequently increased Antarctic glaciation^34^. Furthermore, our analysis highlights that many speciation events in each major family took place within the last 5 million years (**Fig. 2A**). This puts the notothenioid radiation in a similar temporal context as the massive adaptive radiation of cichlid fishes in the African Lake Tanganyika^35^, which also diversified rapidly in that timeframe.

**Figure 2.**
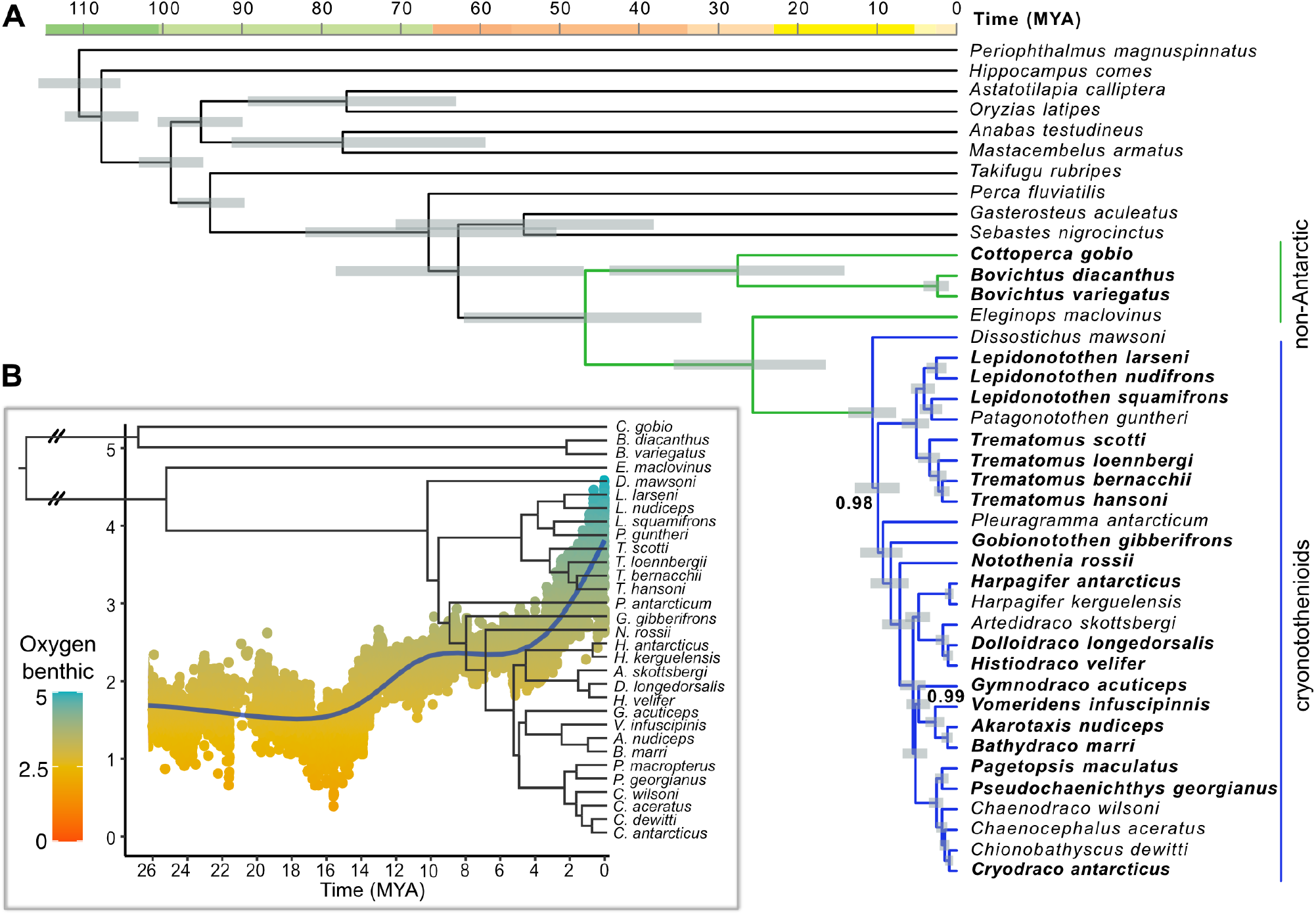
New time calibrated phylogeny and paleoclimate analysis. A) Time calibrated phylogeny of 41 percomorph fish species, including 31 species of notothenioids and 10 outgroup fish species, generated with BEAST2. Branch length corresponds to time in million years (MYA) and grey rectangles show 95% highest posterior density intervals for node age estimates. All nodes received full support (Bayesian posterior probability = 1) except where noted. Species in bold were sequenced in this study. Branches for the Antarctic clade are highlighted in blue (cryonotothenioids), and non-Antarctic notothenioid species are marked in green. B) Diversification of notothenioid species and temperature variation through time. Tree based on notothenioid species from panel A. The scatterplot shows data based on deep-sea δ^18^O benthic records which inversely reflect temperature with higher δ^18^O benthic corresponding to lower temperatures^36^; blue line shows moving average (Generalised Additive Model).

To investigate climatic events that might have driven diversification in derived notothenioid clades we examined paleoclimate data, represented by δ^18^O records (derived from the ratio of ^18^O/^16^O stable isotopes), which reflect variations in the temperature of seawater^36^. These data indicate substantial fluctuations in global mean sea levels (GMSL) during the early Pliocene, and a sustained temperature drop in Antarctica approximately 3 MYA, which led to the rapid formation of large sea ice sheets^36^. Variations in sea ice formation may have played an important role in isolating populations, leading to further diversification of the cryonotothenioids (**Fig. 2B**). A similar influence of cooling events has been suggested in other non-Antarctic radiations^37,38^, where diversification has been linked to past changes in global temperature.

Furthermore, our BEAST2 analyses support the monophyly of dragonfishes (family Bathydraconidae, represented here by *Vomeridens infuscipinnis, Akarotaxis nudiceps, Bathydraco marri*, and *G. acuticeps*),as indicated by morphology and by RADseq data^29^, while other methods and previous studies^30,39^ suggested that they are paraphyletic (**Supplementary Results**). In contrast to previous studies^29,40^, our analyses do not group together the neutrally buoyant *Pleuragramma antarcticum* and *Dissostichus* spp. We therefore suggest that neutral buoyancy evolved independently in the two lineages.

### Transposon expansion is driving genome size evolution in notothenioids

Transposable element dynamics are increasingly recognised as major drivers of evolutionary innovation, and their analysis is greatly facilitated by the use of long-read sequencing technologies^41,42^. For example, the location of TE insertions can influence the expression of nearby genes and induce phenotypic variation^43,44^. The diversity of transposons varies substantially between organisms, with teleost fish genomes containing greater TE diversity compared to other vertebrates, such as mammals^41,45^. In teleosts, genome size tends to correlate with transposon abundance, while a reduction in genome size does not necessarily correspond to lower transposon diversity, but is more commonly caused by reduced copy numbers of TEs^46,47^. Here, we use a set of long read and linked read assemblies, together with high quality *de novo* annotations, to investigate the expansions of transposable elements in notothenioids in relation to their genome sizes. We also investigate the timing of these expansions with respect to major lineage diversification events in the radiation.

We identified substantial variation in assembled genome size across the notothenioid phylogeny with the smallest genomes identified in the basal non-Antarctic temperate-water family Bovichtidae (0.6 Gb) and the largest genomes in the high-latitude icefish species of the derived family Channichthyidae (1.1 Gb) (**Fig. 3C, Extended Data Table 1, Supplementary Table ST2**). This observation is consistent with earlier estimates of large genome sizes in icefish based on flow cytometry^48^. The variation in genome size is essentially completely accounted for by changes in the total repeat content, suggesting that it is driven by TE expansions (**Fig. 3C**). Such expansions are found in diverse members of the Antarctic cryonotothenioids, including *D. mawsoni*, the sister lineage to all other cryonotothenioids, indicating that the onset of TE expansion was associated with the radiation of the clade (**Fig. 3A-B**). This potentially indicates that the onset of the TE expansion coincided with, or possibly predated, the first diversification event in cryonotothenioids. TE expansion continued in the more derived clades (e.g., dragonfishes and icefishes), consistent with lineage-dependent expansion characterised by multiple young insertions as seen in a TE landscape analysis (**Fig. 3B**). Further, we found that this increase in TE content is due to the simultaneous amplification of multiple TE families, including both DNA transposons and retroelements, with the proportion in overall coverage remaining fairly stable throughout the phylogeny (**Fig. 3C, Extended Data Fig. 2, Supplementary Table ST5**). Overall, the bulk of the expansion seems to have resulted from the activation of existing TE families, as several TE families present in the Antarctic clade are also present at low copy numbers in the basal Bovichtidae. Few TE families were observed exclusively within individual clades, although some unclassified TE elements remained in the dataset even after extensive manual curation.

**Figure 3.**
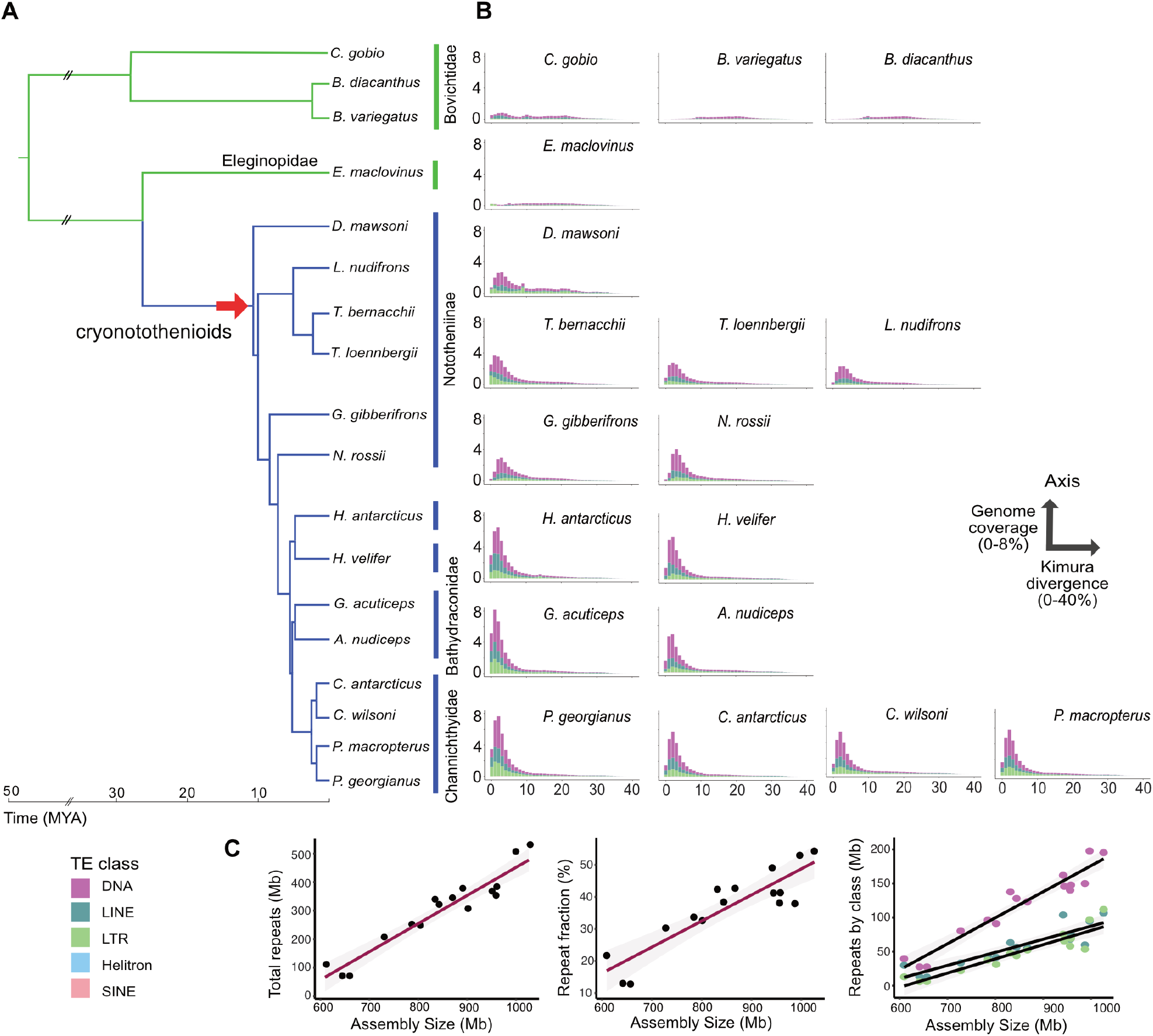
Expansion of transposable elements and genome size variation in notothenioid genomes. A) Species analysed, including 16 species sequenced in this study, and two previously published genomes: *E. maclovinus*^32^, and *D. mawsoni*^54^. B) Repeat landscape plots show distribution of transposable element copies as percentage of divergence from consensus repeat models (x-axis, Kimura divergence) versus genome coverage (y-axis). The red arrow indicates the timing of the earliest TE expansion identified in our analysis. C) Correlation of repeat content with genome size (Pearson Correlation Coefficient, R=0.95, p<0.0001, slope=0.99), increase of repeat fraction with genome size, and increase of DNA, LINE, and LTR TE classes with genome size. Shaded zone indicates 95% confidence interval. Double forward slashes indicate a cropped line in the tree branches and time axis.

In notothenioids, the capacity of transposons to generate evolutionary novelty and shape the evolutionary potential of whole lineages^42^ could be linked to the development of the adaptive features that characterise this radiation. To assess the influence of these transposon events on the genomic evolution of notothenioids, we selected the antifreeze glycoprotein (*afgp*) and the haemoglobin genomic loci as representative models for in-depth examinations.

### Evolution of the antifreeze glycoprotein gene family

The appearance of the antifreeze glycoprotein genes (*afgp*) in Antarctic notothenioid fishes was probably the most important innovation enabling survival in the sub-zero waters of the Southern Ocean. AFGPs prevent organismal freezing by binding to ice crystals that enter the body, thereby arresting ice growth^49,50^. The multigene *afgp* family encodes an array of AFGP size isoforms, whereby each gene is composed of two exons, exon 1 (E1) encoding a signal peptide, and exon 2 (E2) encoding an AFGP polyprotein^10,51^ (**Extended Data Fig. 3A).** The long polyprotein precursor comprises many AFGP molecules composed of varying numbers of repeats of a tripeptide (Thr-Ala-Ala), linked by conserved three-residue spacers (mostly Leu-X-Phe), which on post-translational removal yield the mature AFGPs^10,51,52^. Taken together, the tandemly arrayed *afgp* copies with their extremely repetitive coding sequences present formidable bioinformatic challenges, precluding accurate sequence assemblies and reconstructions of the entire antifreeze gene locus from genomic data until now. The most comprehensive representation of the locus to date was assembled from Sanger-based sequencing of BAC clones for *D. mawsoni*, although this still contains gaps and uncertainties^53^. Furthermore, the evolutionary derivation of *afgp* from its *trypsinogen-like protease* (*tlp*) ancestral gene has not yet been fully resolved. Other uncertainties include the persistence of putative evolutionary intermediates, which are the chimeric *afgp/tlp* genes in extant members of the cryonotothenioids, the origin and the mode by which the three-residue linker sequence appeared, and finally the mechanism of expansion of *afgp* genes^53^.

Using long read data, we assembled the entire *afgp* locus into a single contiguous genomic sequence for a derived icefish species (*P. georgianus).* We also assembled the region corresponding to this locus in a species from a clade that separated prior to the appearance of *afgps (C. gobio*) (**Fig. 4A1 and A3**). For comparison, we reanalysed the *afgp* locus of *D. mawsoni* (**Fig. 4A2**), which represents the most basal cryonotothenioid lineage that diverged after *afgp* emergence^53^. Additionally, we annotated *afgp* genes in three more genomes (*H. antarcticus, T. bernacchii, G. acuticeps*) and located them in multiple scaffolds (**Extended Data Fig. 4**). Manual reassembly to resolve the breaks in these gene clusters of these three species was not possible due to the lack of sufficiently rich long-range data. The assembly of the *afgp* locus in *P. georgianus*, which spans more than 1 Mb in length (1,074 kb), was manually curated to correct mis-assemblies and to verify gene completeness (**Methods**). This *P. georgianus* locus is approximately ten times the length of the corresponding region (113 kb) in *C. gobio*, and more than twice the length of the intermediate *D. mawsoni* locus (467 kb). Overall we observe a remarkable evolutionary expansion that appears to have accelerated in icefishes (**Fig. 4A3**).

**Figure 4.**
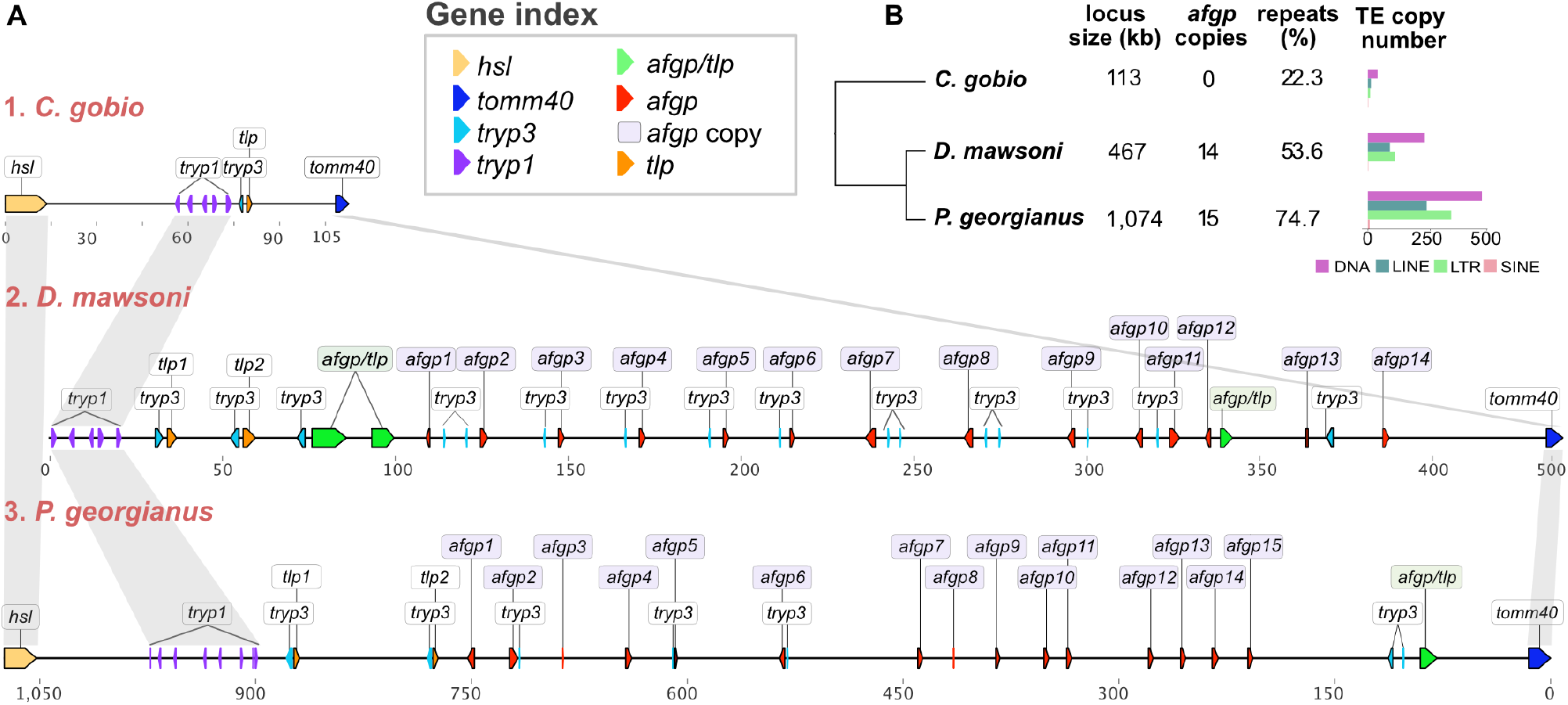
Reconstruction of the antifreeze locus. A) Reconstructed physical map of the antifreeze locus for three notothenioid species: 1) *C. gobio*, which represents the ancestral state of the locus, 2) *D. mawsoni*, and 3) *P. georgianus*, which represent derived loci. *afgp:* antifreeze glycoprotein genes, *tlp*: trypsinogen-like protease, *tryp1:* trypsinogen1, *tryp3:* trypsinogen3 (both *tryp1* and *tryp3* are *prss59* homologues), *tomm40:* translocase of outer mitochondrial membrane 40 homolog, *hsl:* hormone sensitive lipase (*lipeb), afgp/tlp:* chimeric *afgp* and *tlp* gene. B) Cladogram of the three species analysed, with total length of locus, repeat content (%), number of *afgp* gene copies, and number of transposon copies annotated in each species locus (including DNA, LINE, LTRs, and SINE elements). The *C. gobio* and *D. mawsoni* loci are shown at the same scale, and the *P. georgianus* locus is shown in half scale and reverse orientation.

We identified 15 *afgp* genes in *P. georgianus* (**Fig. 4A3**), of which eight are canonical in structure, while the other seven contain various mutations and are thus potentially pseudogenes. In addition there is one chimeric *afgp/tlp* gene, regarded as a putative evolutionary intermediate of *afgp* genes. This also appears to be a pseudogene, because it lacks the signal peptide exon-1, possesses a premature termination codon in the AFGP-coding exon-2, and exon-2 would encode a single long run of 722 Thr-Ala-Ala tripeptide repeats (~6.5 kb) without any of the conserved, cleavable tripeptide linker sequences of the canonical *afgp* polyprotein genes (**Extended Data Fig. 3B)**. Re-annotation of the *D. mawsoni afgp* region identified 14 *afgp* copies, and three chimeric *afgp/tlp* genes, one of which appears to be canonical and potentially functional (**Fig. 4A2**).

The large size discrepancy between the *P. georgianus* and *D. mawsoni afgp* locus cannot be explained by the expansion of *afgp* genes alone, as only one extra copy was found in the icefish species, but instead seems to be primarily due to an expansion of TEs. The repeat content of the locus substantially exceeds the average TE content of the respective genomes (*D. mawsoni:* 53.6% compared to 40.1% genome average; *P. georgianus:* 74.6% compared to 54.3%), consistent with a bias towards TE insertion or retention. We further found evidence of multiple new TE insertions in the region that include representatives of seven LTR, six LINE, and 18 DNA TE families, as well as large expansions of Gypsy, L2, hAT-ac, and Kolobok-T2 copy numbers (**Fig. 4B, Supplementary Table, ST6 Extended Data Fig. 5**). Hence, in addition to transposon copy number being increased by segmental tandem duplication with the *afgp* genes, there has also been active transposition into this region, with the new TE copies potentially involved in local rearrangements.

The physical map of the *afgp* locus presented here for *P. georgianus* represents the most complete reconstruction to date for any icefish species (**Fig. 4**). Previous attempts to map the locus using short read generated assemblies^31^ identified only four to eight copies of the antifreeze genes in various notothenioid genomes. A long-read assembly of *Chaenocephalus aceratus* (blackfin icefish) revealed 11 *afgps*^15^. The locus is also found to be highly fragmented in a recent chromosome-level genome assembly of *D. mawsoni*^54^, once more demonstrating the challenge of assembling this region. Our assemblies also include finer mapping of colocalised, and potentially co-evolving, gene families such as the trypsinogen genes, and tracking of the evolution of the chimeric intermediate gene (*afgp/tlp).* Finally, we observed an inversion of the locus in the icefish in comparison to the *D. mawsoni* haplotype 1 reconstruction^53^.

Two important points emerge regarding the evolutionary progression of the *afgp* gene locus. First, the presence of varying numbers of canonical as well as multiple pseudogene copies of *afgp* genes indicates that the locus remains evolutionarily dynamic. We suggest that maintenance of functional copies in a protein gene family is driven by the strength of selective pressures exerted by the environment. *P. georgianus* is distributed in the considerably milder lower latitudes of the Southern Ocean, around the North of the West Antarctic Peninsula and the Scotia Arc Islands^55^ (**Fig. 1B**), but has never been found in the colder high-latitude waters. Thus, degeneration of previously functional copies would be consistent with a relaxation of selection on maintaining a large functional copy number and energetically costly high level of protein production. Second, the chimeric *afgp/tlp* gene, which was earlier considered to be an evolutionary intermediate state of the early *afgp* genes, can still be found in the icefish. Whilst one chimeric copy remains, it has sustained premature protein truncation and mutation (**Extended Data Fig. 3B**). This is in contrast to the presence of multiple and apparently functional chimeric copies in *D. mawsoni*^53^. Nevertheless, this chimeric gene in *P. georgianus* is clearly a pseudogene on its way to extinction. First, it lacks the signal peptide coding sequence, thus could not produce a secreted protein even if functional. Second, the very long run of coding sequence of *afgp* tripeptide repeats (722 repeats; ~6.5 kb) without any of the conserved cleavable 3-residue linker sequences in canonical *afgp* polyprotein genes is indistinguishable from simple sequence repeats of nine nucleotides. This suggests that once independent functional *afgp* copies were formed and with independent *tlp* already present, the maintenance of a chimeric copy may have become unnecessary in the less selective low latitude habitat ranges that *P. georgianus* colonised.

The evolutionarily dynamic nature of the notothenioid *afgp* gene family can also be gleaned from the sequence of *T. bernacchii*, which is a species that resides in the most severe conditions at the southernmost limit (McMurdo Sound, 78°S) for marine life in the Southern Ocean. Even though its *afgp* locus assembly lacks contiguity, its annotation presents the largest set of *afgp* gene copies of all analysed species (24 copies, of which at least 11 are canonical), and also maintains three chimeric genes (**Extended Data Fig. 4**). Future efforts in assembling the challenging *afgp* loci to contiguity for cryonotothenioids across latitudinal clines will inform on the evolutionary dynamics of the adaptive *afgp* trait as driven by environmental selective strength.

### Gene gains and losses drive haemoglobin evolution in notothenioids

The haemoglobin gene family has also been under strong selective pressure in notothenioids^56^. Haemoglobin is essential for oxygen transport, and the evolution of haemoglobin genes has been fuelled by duplications that enabled diversification of paralogous genes, as well as adaptation to changing environments through alterations in expression patterns^57,58^. In teleosts, haemoglobins are organised in two clusters, each containing both α and β globin genes: the larger MN cluster (flanked by genes *mpg* and *nprl3*), and the smaller LA cluster (flanked by *lcmt1* and *aqp8*)^59^. Relaxed selection on haemoglobins and red blood cells in the cold, oxygen-rich Southern Ocean has led to moderate to severe anaemia in multiple notothenioid lineages^60^. The icefish family (Channichthyidae), often called “white blooded” icefishes due to their translucent white blood, are the only known vertebrates that completely lack haemoglobin and do not produce mature erythrocytes. Instead, they rely on oxygen physically dissolved in the blood plasma, which is possible because of the high oxygen saturation level at the freezing temperatures of the Southern Ocean^61^. The mechanisms underlying the loss of haemoglobin genes in the icefish are still a mystery, partly because the reconstruction of complete haemoglobin loci was not possible with previous fragmented genome assemblies. Here we describe the most complete reconstruction of the haemoglobin loci in notothenioids to date, allowing us to evaluate their evolution, track the loss of haemoglobins in the icefish, and identify potential involvement of transposable elements in this process.

Using five new long-read assemblies, we achieved the contiguous assembly of both LA and MN haemoglobin gene clusters for most of the species, including their flanking genes^59,62^, and compared these with four published assemblies for three notothenioids (*Eleginops maclovinus*^32^, *D. mawsoni*^54^, and *C. aceratus*^15^) and one temperate non-notothenioid perciform (*Perca flavescens*^63^) (**Fig. 5, Extended Data Fig. 6**). The two loci were found to be distinct^64^ and located on two different chromosomes (LA: chr19, MN: chr8) that originated in the teleost genome duplication^65^. We devised and use here a new naming system for teleost haemoglobin genes independent of the life stage of expression (**Supplementary Methods**). In the icefish species examined, we confirm the complete loss of functional haemoglobin genes in both clusters, with a single pseudogenized gene copy remaining in the LA locus. Synteny analysis suggests that the loss of haemoglobins potentially occurred as a single event for each locus. Furthermore, we find that the remaining length of the MN locus in *P. georgianus* is characterised by multiple transposon insertions (**Fig. 5B**). No other coding genes are found in the region, and the total length of the remaining genomic region is approximately the same size as that of red-blooded cryonotothenioids (30-60 kb). The most common transposon insertions include LINE/L2 elements, which account for approximately 20% of the total length of the *P. georgianus* MN locus (**Supplementary Table ST7**, **Extended Data Fig. 7)**. Similar transposon insertions are found in the LA locus, where the only haemoglobin remnant is the third exon of the pseudogenized *α-globin.2* (**Fig. 5B**), as previously identified^15,64,66^.

**Figure 5.**
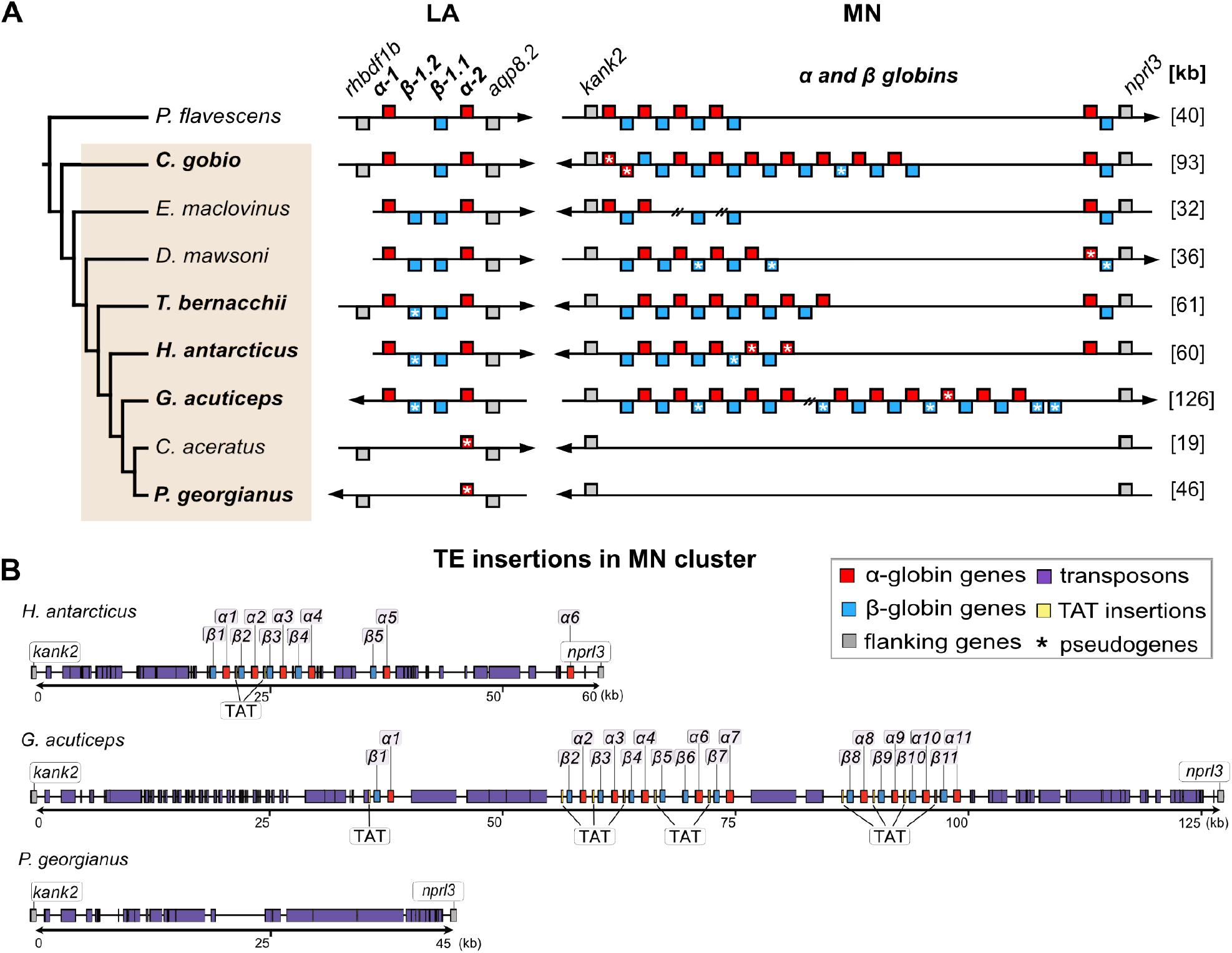
Reconstructed synteny of haemoglobin loci and TE insertions. A) Species analysed and syntenic reconstruction of LA and MN haemoglobin gene clusters. B) Transposon insertions in the MN region of *H. antarcticus, G. acuticeps*, and *P. georgianus* genomes. Red: *α-globin* genes, blue: *β-globin* genes, grey: flanking genes, purple: transposon insertions, yellow: TAT-like repeats. Pseudogenes are marked with asterisks. Breaks in the assembly are indicated with double forward slashes. Bold face indicates species sequenced in the present study. Arrows show locus orientation and total lengths of MN locus in different species are given in brackets (kb, at right).

In contrast, in red-blooded species, haemoglobin loci are characterised by several local duplications (**Fig. 5**). For the LA cluster, in the common ancestor of *E. maclovinus* and cryonotothenioids we find a previously unidentified duplication of the *β-globin.1* gene that gave rise to two β-globin copies (*β-globin1.1* and *1.2*) (**Fig. 5A**). Subsequently, one duplicated copy was repeatedly lost or deleteriously altered in other cryonotothenioid species, while the other copy was retained throughout, until completely lost in the icefish. In the larger MN cluster, the number of haemoglobin gene copies varies considerably across species, with five to 11 α-globin and five to 13 β-globin genes, while containing multiple lineage-specific tandem duplications of the *α-globin.1* and *β-globin.1* gene pair in every lineage analysed (α-globin and β-globin genes, **Fig. 5A**). We retrieved a lower number of copies in the MN locus of the one species for which short read data were used for assembly (*E. maclovinus*^32^), potentially due to incomplete assembly or misassembly. Several cryonotothenioid species display lineage-specific frame-shifts or premature stop codons predicted to cause loss of function. The nature of the pseudogenisation of these gene copies suggests multiple independent pseudogenisation events. Only the loss of the first *α-globin.1* copy appears to be shared across cryonotothenioids. At the opposite end of the cluster, the *α-globin.2* and *β-globin.2* pair was retained with, however, the notable loss of *β-globin.2* in *H. antarcticus*, and in *G. acuticeps.* The case of *G. acuticeps* is particularly interesting, as here we identify a massive expansion of haemoglobin genes in the MN cluster, comprising more than twice as many haemoglobin genes as in other cryonotothenioid species. *G. acuticeps* is a member of the Bathydraconidae (dragonfishes), the group most closely related to the haemoglobin-lacking icefish. Furthermore, we find that each haemoglobin gene pair is preceded by a TAT-like repeat insertion, consistent with non-homologous recombination between these repeats underlying the tandem expansion of gene pairs (**Fig. 5B**). In other globin genes, such as myoglobins, TAT-like insertions have been shown to interfere with transcription regulation^67,68^. Further analysis of haemoglobin expression levels would be needed to understand how these TAT-like insertions influence haemoglobin transcription regulation in notothenioids.

By assembling the most complete reconstruction of notothenioid haemoglobin loci to date, we track the patterns of loss and gain of haemoglobin copies across the radiation and identify transposon insertions that may have influenced their evolution, while syntenic analysis suggests that the loss of haemoglobins in icefish could be potentially attributed to a single deletion event. Though it has been suggested that the lack of erythrocytes may reduce blood viscosity, the energy required to pump high volumes of blood and the need for additional physiological adaptations, puts into question whether the loss of haemoglobins in icefishes is indeed an adaptive trait^56^. Perhaps the lack of intense niche competition aided the establishment of this phenotype in the highly oxygenated waters of the Southern Ocean in the past. However, relying on an oxygen absorption system that depends on oxygen diffusion leaves the icefish intensely vulnerable to rising temperatures and thus decreased dissolved oxygen in the future^61^.

## Conclusions

We present the most extensive effort to date to investigate the genomic evolution underlying the iconic notothenioid fish radiation, via the generation of a set of 24 new genome assemblies encompassing representatives from almost all notothenioid families. We demonstrate that the use of high-quality genome assemblies over a wide taxonomic breadth can help to decipher the evolutionary history of this recent vertebrate radiation. We identify critical steps in the evolution of key gene families that involve large genomic rearrangements in repetitive regions that could only be reconstructed with the aid of long-read assemblies. We show that the evolutionary history of the remarkable notothenioid radiation was associated with, and potentially driven by, transposon proliferation, which could have affected the evolutionary advantage of these species during freezing events occurring in the Southern Ocean. In particular, TE expansion events can be linked to the structure of characteristic repetitive gene families such as the haemoglobin and antifreeze genes. Beyond these direct insights, our work provides an extensive resource for the future study of notothenioid genomic evolution, enabling further research to advance our understanding of the notothenioid radiation, and of genomic adaptations to extreme environments more widely. As the fate of notothenioid diversity is linked to very narrow margins of temperature tolerance, studying their adaptability is particularly relevant now with the unfolding climate crisis and the warming of the Southern Ocean^69^.

## Methods

### Sample processing and sequencing

Tissue samples were collected and either frozen immediately at −80 °C, or preserved in ethanol and then frozen to preserve the quality of genomic DNA. Tissue preservation can influence the quality of extracted DNA^70^, and flash freezing is the optimal preservation method for use with long-read sequencing. High molecular weight DNA (HMW DNA) was extracted using the Bionano Agarose plug extraction protocol^71^ or a modified version of the MagAttract kit (Qiagen) (**Supplementary Table ST1**). The quantity of extracted HMW DNA was evaluated with the HS Qubit DNA kit and the fragment profile and overall quality was assessed with the Femto Pulse instrument (Agilent). Pacific Biosciences (PacBio) sequencing was performed with CLR (Continuous Long Reads) SMRT cells. PacBio libraries were made using the SMRTbell Template Prep Kit 1.0, following the PacBio protocol. A size selection was performed on the BluePippin instrument (Sage Science), with a 15kb cut off and sequence data were generated on the Sequel instrument using Seq kit v2/Binding Kit v2.0 with a 10 hour movie. Illumina sequencing paired end (PE) libraries were generated for eight species and sequenced by multiplexing two species per lane on Illumina HiSeqX (150 bp PE). Linked reads 10X Genomics Chromium sequencing was performed for 17 species (**Extended Data Table 1**). Linked-read libraries were prepared using the Chromium Genome Reagent Kit, and the Chromium Genome Library Kit & Gel Bead Kit, according to the manufacturer’s instructions^72,73^, with standard DNA input (1 ng). Hi-C libraries were generated with a Dovetail kit for *P. georgianus* and with ARIMA Genomics kit for *C. gobio*, and each was sequenced on the Illumina HiSeq4000 platform. Bionano Irys optical mapping was used for scaffolding the assembly of *H. antarcticus*, as well as that of *C. gobio*^23^. Total RNA for RNAseq was extracted using the RNeasy Qiagen extraction kit, and Illumina 150 bp PE libraries were sequenced on the HiSeq4000 platform (**Supplementary Methods**). All sample processing and sequencing was performed at the Wellcome Sanger Institute, UK.

### Genome assembly and curation

For genome assembly we used a combination of different sequencing technologies, which were either used in conjunction (hybrid assemblies) or individually. Detailed information on the assembly process used for each species are given in the **Supplementary Methods**. Briefly, our genome assemblies were generated as follows.

For *C. gobio* the assembly was generated based on 75x PacBio Sequel data, 54x Illumina HiSeqX data generated from a 10X Genomics Chromium library, Bionano Saphyr two-enzyme data (Irys) and 145x coverage HiSeqX data from a Hi-C library (for Hi-C, tissue from a different individual was used), as described in ref^23^. For *P. georgianus* the assembly was based on 93x PacBio, 56x 10X Genomics Chromium, and Dovetail Hi-C data, and was generated using Falcon-unzip with 10X based scaffolding with scaff10x, BioNano hybrid-scaffolding, Hi-C based scaffolding with SALSA2^74^, Arrow polishing, and two rounds of FreeBayes^75^ polishing. This assembly comprises 24 chromosomes, which were numbered in correspondence to the medaka HdR1 assembly (*Oryzias latipes*, GCA_002234675.1), as for the *C. gobio* genome^23^. The assembly for *H. antarcticus* was based on 67x PacBio data, 40x of 10X Genomics Chromium data, and Bionano Irys data. The initial PacBio assembly was generated with Falcon-unzip, retained haplotig identification with Purge Haplotigs^76^, 10X based scaffolding with scaff10x, Bionano hybrid-scaffolding, Arrow polishing, and two rounds of FreeBayes polishing. For *T. bernacchii* and *G. acuticeps* the assemblies were based on PacBio and 10X data. For *T. bernacchii*, we used 46x PacBio data and 53x of 10X Genomics Chromium data, while the assembly for *G. acuticeps* was based on 31x PacBio data and 41.8x of 10X Genomics Chromium data. These were assembled using Falcon-unzip, 10X based scaffolding with scaff10x, Arrow polishing, and two rounds of FreeBayes polishing. Purge_dups^77^ was run on the curated assemblies to further remove retained duplications.

For 19 more species only 10X Chromium or Illumina HiSeqX were used for assembly. (**Extended Data Table 1**). For 11 species sequenced with 10X Genomics Chromium data, genome assembly was performed using Supernova 2.0 (<u>Supernova 2.0 Software</u>). After initial assembly, retained haplotigs were identified using Purge Haplotigs^76^. For the remaining eight species which were only sequenced with Illumina HiSeqX, a primary assembly was generated with a reference guided approach using SOAPdenovo2^25^. The short insert reads were initially base error corrected using BFC (https://github.com/lh3/bfc). After this step, larger kmer sizes (e.g. 70), may be applied to improve assembly. SOAPdenovo^25^ was used to process the cleaned short insert reads, followed by GapCloser for contig gap filling. The scaffolds were further enhanced by the use of cross_genome, a tool which maps genome synteny to merge scaffolds (<u>Phusion2 - Browse /cross_genome</u>), as has been previously applied in genome assemblies such as Tasmanian devil^78^ and grass carp^79^. To finalise these assemblies, decontamination methods were used to remove contaminants from sequencing (e.g. adapter sequences) or symbionts. Metrics for sequencing data and assemblies are provided in **Supplementary Tables ST8 and ST9**.

To improve the quality of the PacBio assemblies we performed manual curation to remove mis-assemblies, duplications, and sequencing contamination, and to merge scaffolds based on supporting evidence, which has been shown to substantially improve the continuity and accuracy of genome assemblies^20^. Each assembly was manually curated using the Genome Evaluation Browser (gEVAL)^19^. Scaffold integrity was confirmed with PacBio and 10X illumina read mapping, while for *H. antarcticus* scaffold integrity was further confirmed using Bionano BssSI optical maps. For *P. georgianus* a 2D map was built using Hi-C reads, allowing further scaffold correction and super-scaffolding to bring the assembly to chromosome scale. Artificially retained haplotypic duplications were removed with purge_dups^77^. Chromosome name assignment was based on comparative alignment to the *C. gobio* chromosome assembly (GCA_900634415.1) (**Supplementary Methods, Supplementary Table ST10**).

### Gene and transposable element annotation

Gene annotation was generated by Ensembl for all five PacBio assemblies, using the Ensembl Gene Annotation system^27^. For *C. gobio* assembly (fCotGob3.1), transcriptomic data were generated for four tissue types (brain, muscle, gonad, and spleen). We also sequenced for *T. bernacchii* brain, muscle, and gonads (male and female), and brain and muscle for *G. acuticeps.* Annotations are available on the Ensembl server under GCA_900634415.1 (database version 9.31^77^) for *C. gobio*, and on Ensembl Rapid Release for *G. acuticeps, P. georgianus, H. antarcticus*, and *T. bernacchii*, (**Supplementary Methods, Supplementary Table ST4**). Comparison of orthologous clusters was performed with OrthoVenn2^80^.

To calculate overall repeat content for each species we used WindowMasker v.1.0.0^81^, a method that does not require a repeat library. *De novo* annotation of transposable elements was performed using RepeatModeler v.2.0^82^ and RepeatMasker v.4.0.1^83^. For each of the five PacBio assemblies (*C. gobio, T. bernacchii, H. antarcticus, G. acuticeps*, and *P. georgianus*) a *de novo* repeat library was generated using RepeatModeler2^82^. To enhance detection of LTR retrotransposons, the programs LTRharvest^84^ and LTR_retriever^85^ were run as part of the RepeatModeler2 pipeline. The identified elements from each genome were further curated manually to remove false assignments and achieve complete length elements. The consensus TE sequences identified by RepeatModeler2 were blasted against the genomes (BLAST+^86^), and the sequences of the 50 best hits were extracted along with a 1-kb long flanking sequence on each side. Multiple sequence alignments were generated for each set of top hits, using MUSCLE^87^. Each multiple sequence alignment was visualised with belvu^88^, and then manually inspected to confirm TE element completeness. TEs that appeared to be extending beyond alignment boundaries were subjected to additional rounds of curation, until the complete sequence was recovered. Finally, new consensus sequences were extracted from the multiple alignments using hmmer (http://hmmer.org/). Overall a total of ~2,000 elements generated by RepeatModeler2 were manually curated. A custom TE library was created combined with the curated elements and LTR outputs and all genomes were masked with RepeatMasker. TE copy numbers were calculated using a Perl script parsing the RepeatMasker output^89^.

### Phylogenetic analysis

Phylogenetic analysis was performed using single copy ortholog genes identified with BUSCO^26^, for the 24 newly sequenced notothenioid genomes and 17 previously published genomes of seven notothenioids and ten further species of percomorph fishes. The species and assembly versions used are listed in **Supplementary Table ST11.** BUSCO (v2) was run with lineage “actinopterygii_odb9”, and the sequences of single copy orthologs identified in each assembly and extracted for use in further analysis.

Detailed information about the phylogenetic analyses are provided in the **Supplementary Methods**. Briefly, we used MAFFT v.7.453^90^ to align 266 selected BUSCO genes, and after rigorous manual curation removing alignments with missing data, high entropy, and lengths shorter than 500 bp, 228 alignments remained for further phylogenetic analysis. Each of these alignments was subjected to Bayesian phylogenetic analysis with BEAST2 v.2.6.0^91^, with an uncorrelated lognormal relaxed clock model and a Markov-chain Monte Carlo chain (MCMC) length of 25 million iterations. “Strict” and “permissive” sets of alignments were compiled based on estimates of the mutation rate and its among-species variation, and contained 140 and 200 of the alignments, respectively. For the strict set, the permissive set, and a “full” set (257 alignments; excluding nine unsuitable BUSCO genes), we also performed maximum-likelihood phylogenetic analyses with IQ-TREE v.1.7^92^. Each of the three analyses was complemented with an estimation of gene- and site-specific concordance factors, and the three resulting sets of gene trees were used for separate species-tree analyses with ASTRAL v.5.7.3^93^. Finally, we estimated the phylogeny and the divergence times of notothenioid species with BEAST2 from a concatenated alignment combining all alignments of the strict set. The original data blocks were grouped in 12 positions selected with the rcluster algorithm of PartitionFinder v.2.1.1^94^, assuming linked branch lengths, equal weights for all model parameters, a minimum partition size of 5,000 bp, and the GTR+Gamma substitution model. The same substitution model was also assumed in the BEAST2 analysis, together with the birth-death model of diversification^95^ and the uncorrelated lognormal relaxed clock model^96^. Time calibration of the phylogeny was based on four age constraints defined according to a recent timeline of teleost evolution inferred from genome and fossil information^33^, at the most recent common ancestors of clades: Eupercaria, around 97.47 MYA (2.5–97.5 inter-percentile range: 91.3–104.0 MYA); the clade combining Eupercaria, Ovalentaria, and Anabantaria – around 101.79 MYA (95.4–109.0 MYA); the clade combining these four groups with Syngnatharia and Pelagiaria – around 104.48 MYA (97.3–112.0 MYA); and the clade combining those six groups with Gobiaria – around 107.08 MYA (100.0–114.0 MYA). Additionally, we constrained the unambiguous^33,97–99^ monophyly of the groups Notothenioidei, Perciformes, Ovalentaria, Anabantaria, and the clade combining the latter two groups. We performed six replicate BEAST2 analyses with 330 million MCMC iterations, and convergence among MCMC chains was confirmed for all model parameters. The posterior tree distribution was summarised in the form of a maximum-clade credibility tree with TreeAnnotator v.2.6.0^100^. We attempted to repeat the BEAST2 analyses with the permissive and full datasets, but these proved too computationally demanding to complete. Nevertheless, the preliminary results from these analyses supported the same tree topology as the analyses with the strict dataset.

To place the time calibrated phylogeny in the context of historic ocean temperature variation, we used estimated benthic oxygen values previously published in ref^36^. The moving average was plotted with geom_smooth in R using a generalised additive model (GAM: (y ~ s(x, bs = “cs”)).

### *afgp/tlp* locus annotation and reconstruction

The location of the *afgp* locus in each genome assembly was initially identified with BLAST+^86^ searches, using as query sequence a copy of the *afgp* gene previously sequenced in *D. mawsoni*^53^. Additionally, we searched using sequences from each of five other genes known to flank or colocalize with *afgp* copies, including: *tlp*: trypsinogen-like protease, *tryp1:* trypsinogen1, *tryp3:* trypsinogen3 (both prss59 homologues), *hsl*: hormone sensitive lipase (*lipeb*) and *tomm40* (translocase of outer mitochondrial membrane 40 homolog). The exact location of each gene copy was confirmed and annotated manually, to identify numbers and sizes of exons. Each *afgp* copy was manually inspected for the presence of frame shifts and gaps to identify complete canonical genes and pseudogenized copies.

All five genomes sequenced with PacBio contained copies of all six genes used for BLAST+ analyses (*afgps* and flanking genes) (**Fig. 4**, **Extended Data Fig. 5**). To further improve the assembly of the *afgp* locus on the *P. georgianus* genome, we mapped Falcon-corrected PacBio reads to the diploid assembly using minimap2^101^, and then filtered the mapped reads to remove secondary alignments (samtools view -F 256). We used GAP5^102^ to inspect and manually curate the mapped reads. Reads that mapped to more than one location were linked in GAP5, and by further inspecting these links and extending soft-clipped sequence, it was possible to merge contigs, resulting in a complete representation of the whole *afgp* gene locus. Finally, the reassembled sequence was polished using Racon^103^.

### Haemoglobin gene locus annotation and reconstruction

To study haemoglobin genes in notothenioid species we focused on the five species (*C. gobio, T. bernacchii, H. antarcticus, G. acuticeps*, and *P. georgianus*) assembled with PacBio data, along with three previously published assemblies (*D. mawsoni*^54^, *E. maclovinus*^32^, *and C. aceratus*^15^) to provide as good a clade coverage as possible. The reference genome assembly of the yellow perch (*Perca flavescens*)^63^, a close relative to notothenioids within the order Perciformes^9^, was used as a reference to determine haemoglobin gene exon boundaries. We used flanking genes to confirm orthology between genes and clusters across species, and each exon of each gene was retrieved and their exact positions in the corresponding genome assembly was recorded. Using the *T. bernacchii* assembly as reference species we performed mVISTA^104^ alignments of genomic regions with LAGAN^105^ or Shuffle-LAGAN^106^. Protein sequences were aligned with MUSCLE^87^ and phylogenetic trees were reconstructed with RAxML-NG^107^ using the best-fitting substitution model according to ModelFinder based on Bayesian information criterion (BIC)^108^.

We used self-alignments with dotter^109^ to visualise the complete reconstruction of each haemoglobin cluster and we manually inspected each locus to identify possible gaps or mis-assemblies. We can confirm complete gapless assembly of the MN haemoglobin locus for each PacBio species, with the exception of one gap identified in the *G. acuticeps* genome, which could not be corrected with the available data (**Fig. 5**). Furthermore, the *P. georgianus* chromosomal assembly MN haemoglobin locus was also manually curated using GAP5^102^, as described above for the *afgp* locus.

The previously employed nomenclature system for haemoglobin genes was based on zebrafish species expression patterns (embryonic and adult), but this system may not be transferable across all species^62,110^. Here we have thus employed a novel nomenclature system that incorporates cluster designation information (LA or MN) while relying on genomic organisation rather than expression pattern (**Supplementary Methods**).

Detailed information on all the tools and versions used for each analysis are provided in **Supplementary Table ST12**.

## Extended Data Figures and Tables

**Extended Data Table 1.**
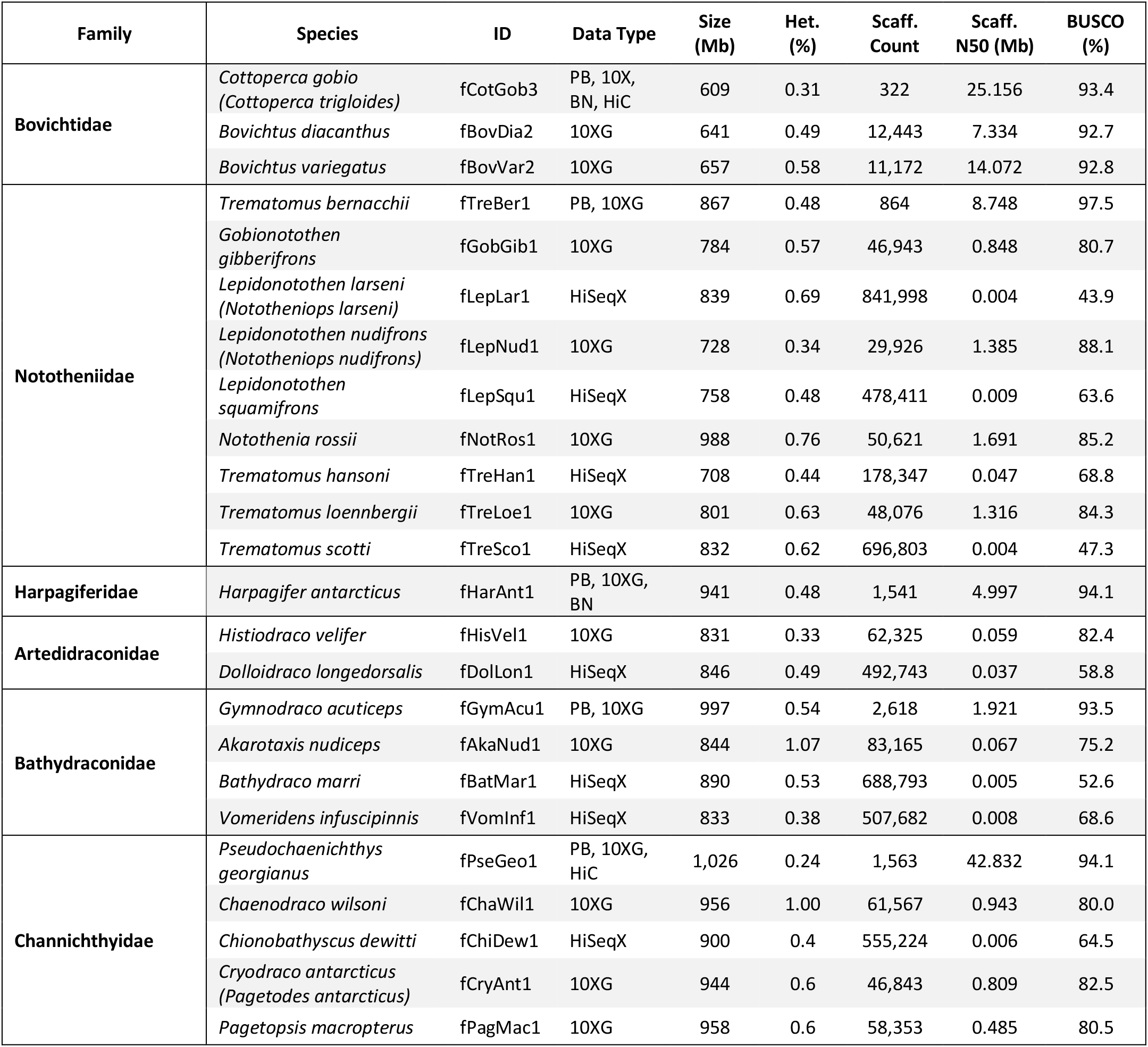
Target species, data types and assembly statistics. Data types used to generate assemblies: PB: PacBio CLR, 10XG: 10X Genomics linked reads, BN: Bionano, HiSeqX: Illumina Hi-Seq, HiC: Hi-C from ARIMA (*C. gobio*), Dovetail (*P. georgianus*). Gene completeness was calculated with BUSCO v.5, and heterozygosity (Het.) estimated with GenomeScope (kmer=31). Names in parentheses are recent nomenclature revisions^3^. Scaff = scaffold.

**Extended Data Figure 1.**
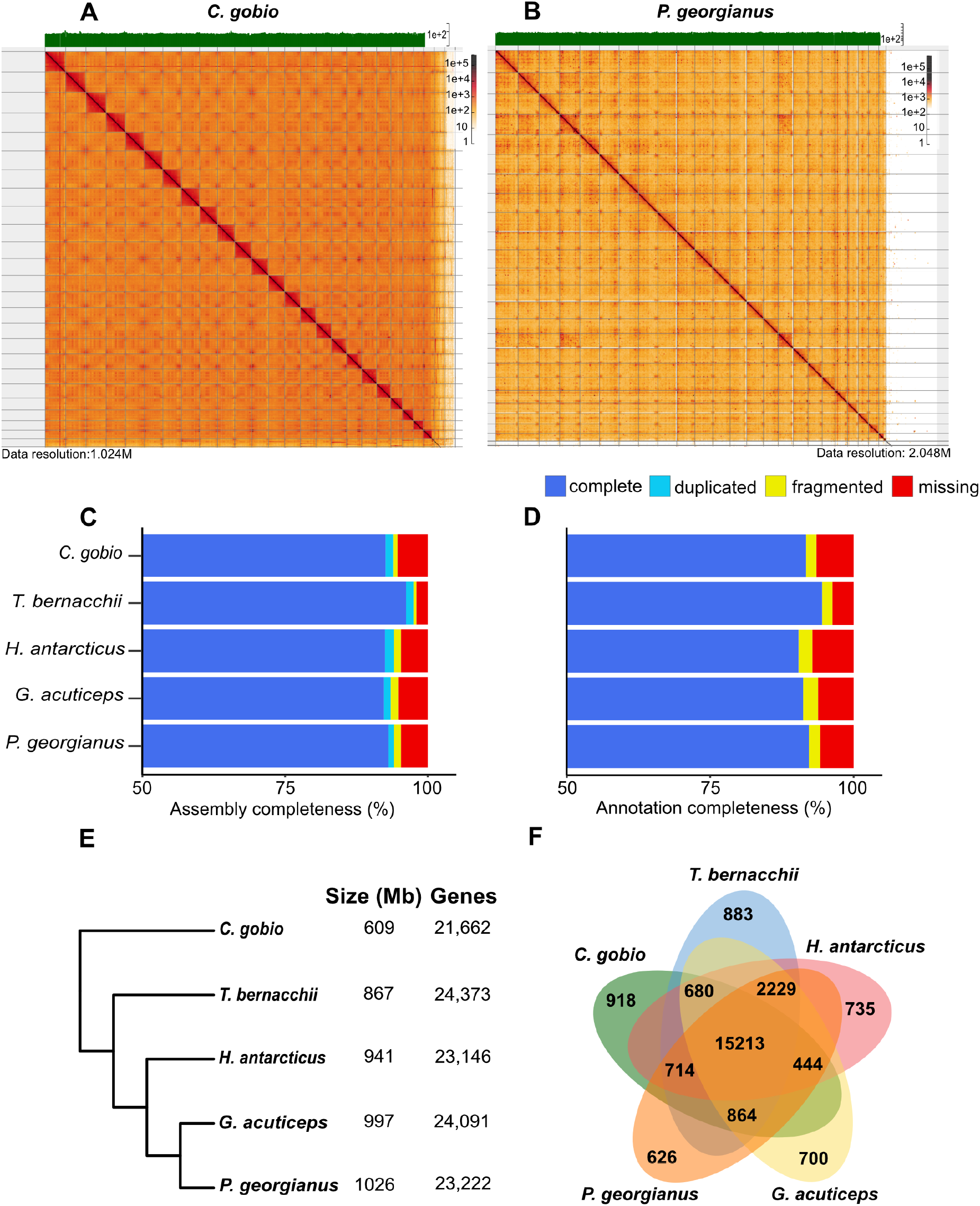
Hi-C maps, assembly and annotation completeness of PacBio genomes. Hi-C maps for species *C. gobio* (A), and *P. georgianus* (B) generated with HighGlass, showing the assemblies scaffolded on 24 chromosomes (red squares across the diagonal), and read coverage (green track on top). C) Assembly and D) annotation completeness scores based on BUSCO (v.5), E) genome size and number of annotated genes (Ensembl) for each PacBio assembly, F) unique and shared Ensembl genes.

**Extended Data Figure 2.**
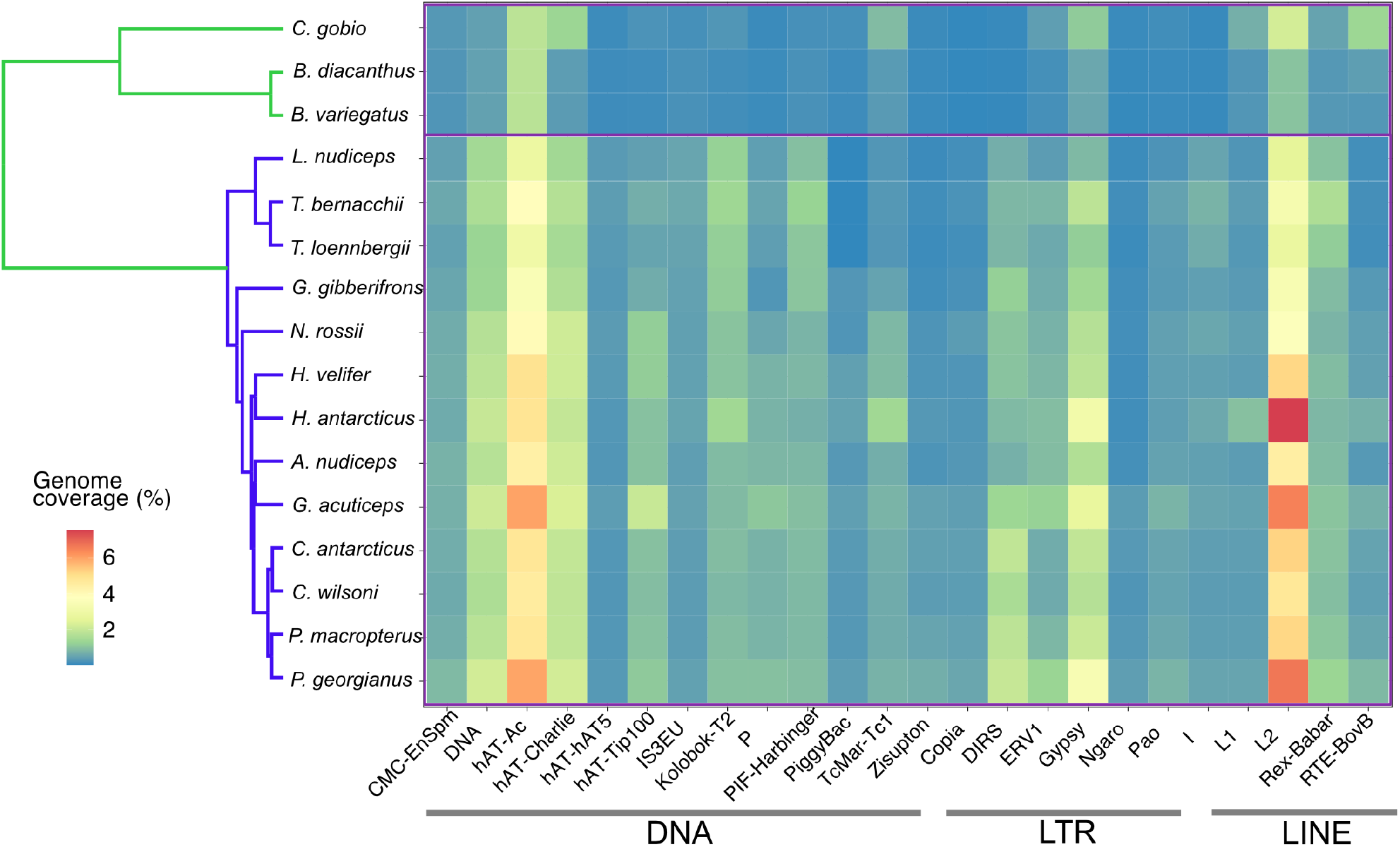
Heatmap of transposon coverage of most abundant transposon superfamilies. Coverage is shown as percent (%) of genome for 16 notothenioid species sequenced in present study, including DNA transposons, and LTR and LINE retrotransposons. The colour of tree branches indicates non-Antarctic (green), and cryonotothenioid (blue) species.

**Extended Data Figure 3.**
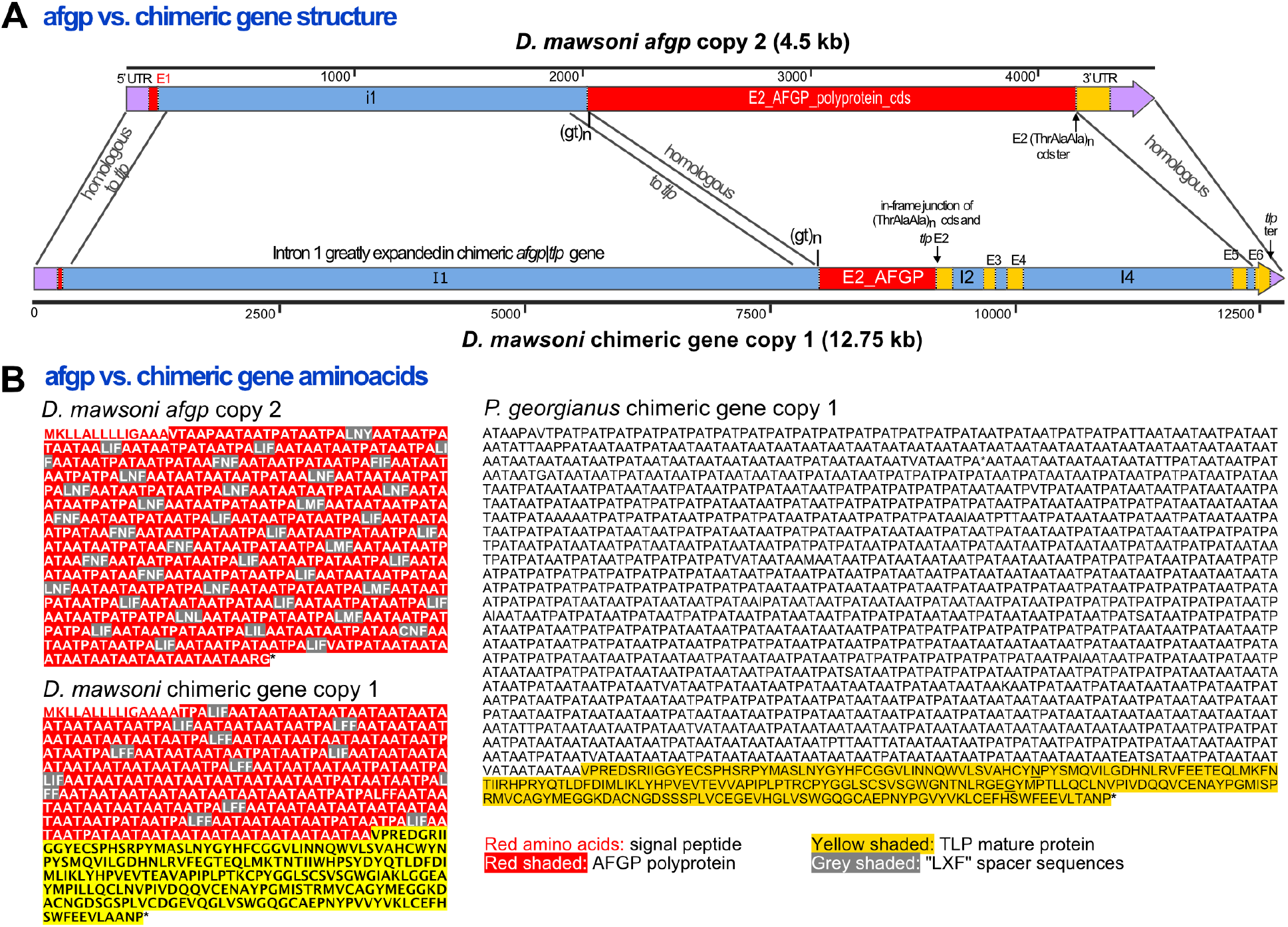
A) Structure of *afgp* gene copy 2 from species *D. mawsoni* showing the two exons (E1: signal peptide, E2: long polyprotein precursor), compared to a chimeric gene (copy 1 of *D. mawsoni*), which contains complete cds of both *afgp* and *tlp* genes. B) Amino acid sequences of *D. mawsoni afgp* copy 2 and chimeric gene copy 1, showing the signal peptide (red letters), AFGP polyprotein chain (red shading), the TLP mature protein chain (yellow shading), and spacer sequences (grey shading). Also shown is the remaining and pseudogenized chimeric gene in *P. georgianus*, which contains neither a signal peptide nor an AFGP polyprotein chain.

**Extended Data Figure 4.**
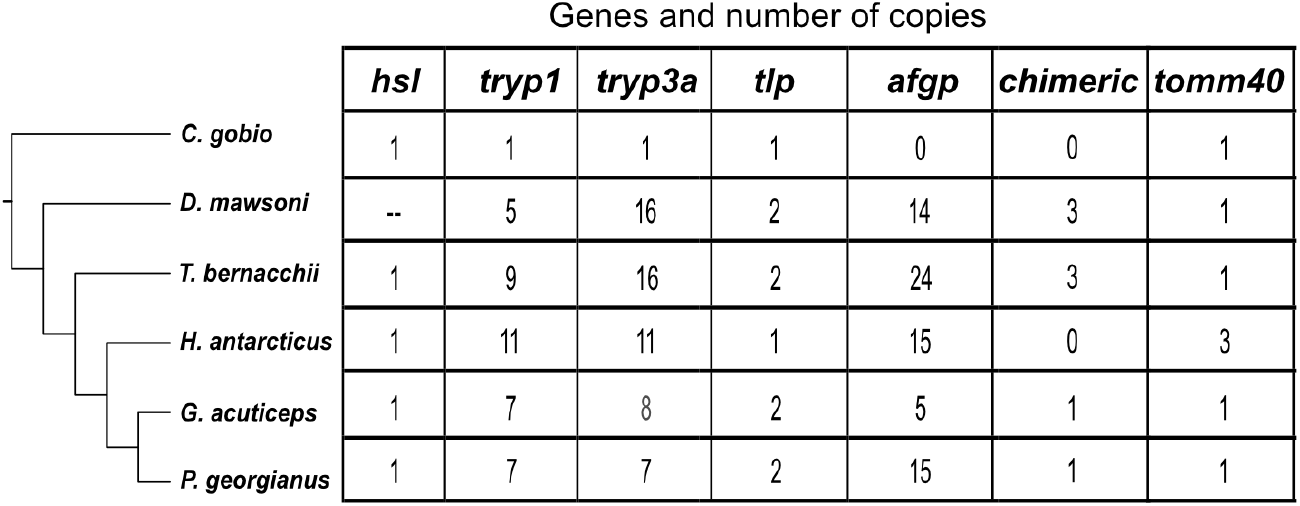
Genes in the *afgp* locus of six notothenioid species and number of copies annotated. Including the following genes: *afgp:* antifreeze glycoprotein genes, *tlp:* trypsinogen-like protease, *tryp1:* trypsinogen1, *tryp3:* trypsinogen3 (both *tryp1* and *tryp3* are *prss59* homologues), *tomm40:* translocase of outer mitochondrial membrane 40 homolog, *hsl:* hormone sensitive lipase (*lipeb), afgp/tlp:* chimeric *afgp* and *tlp* gene.

**Extended Data Figure 5.**
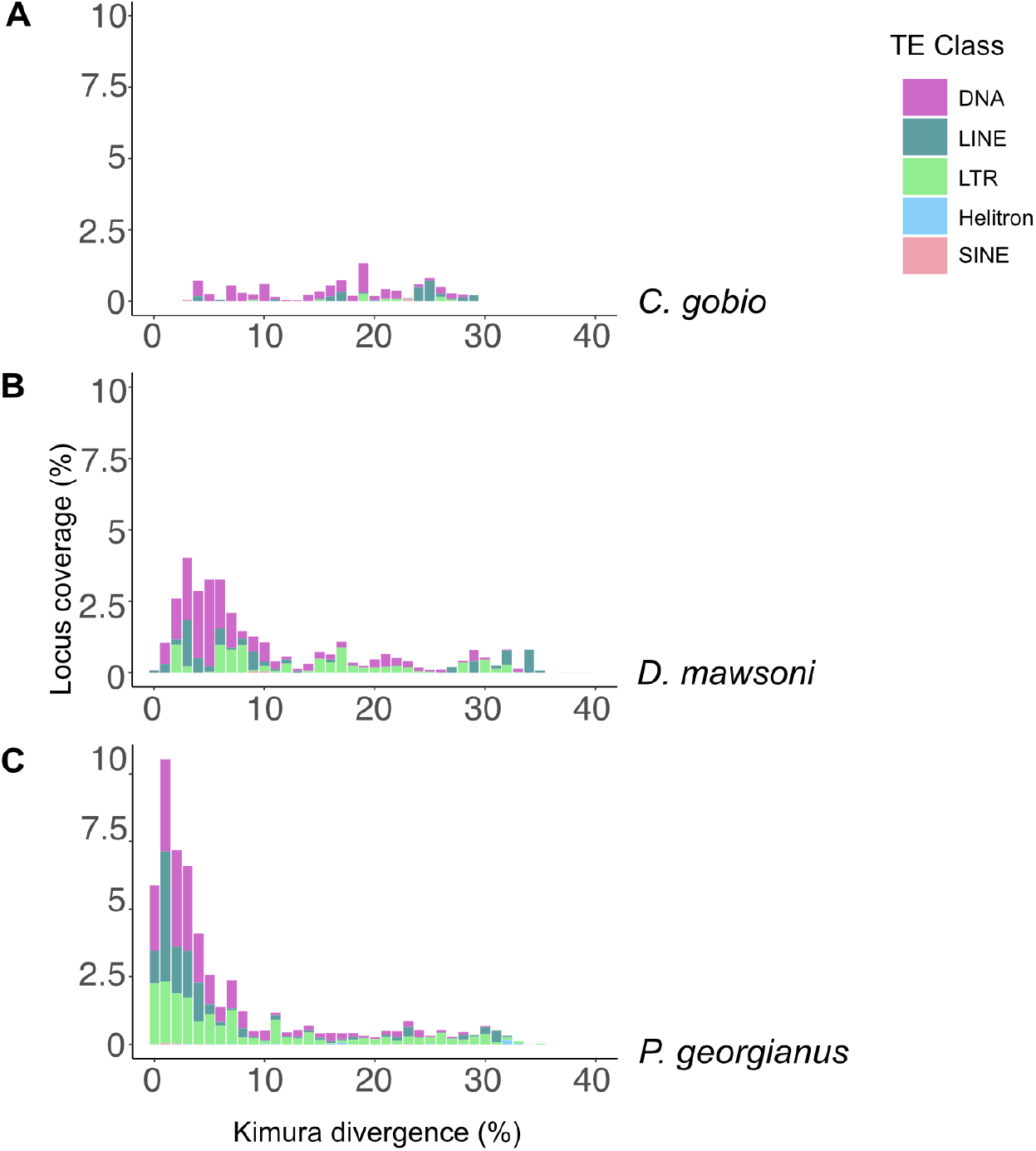
Transposon insertions in the *afgp* locus of three species, A) *C. gobio*, B) *D. mawsoni*, and C) *P. georgianus*. Plots show distribution of TE copies based on % of divergence from their consensus sequences (x-axis), and overall coverage as % of genome (y-axis). The colours represent different TE classes.

**Extended Data Figure 6.**
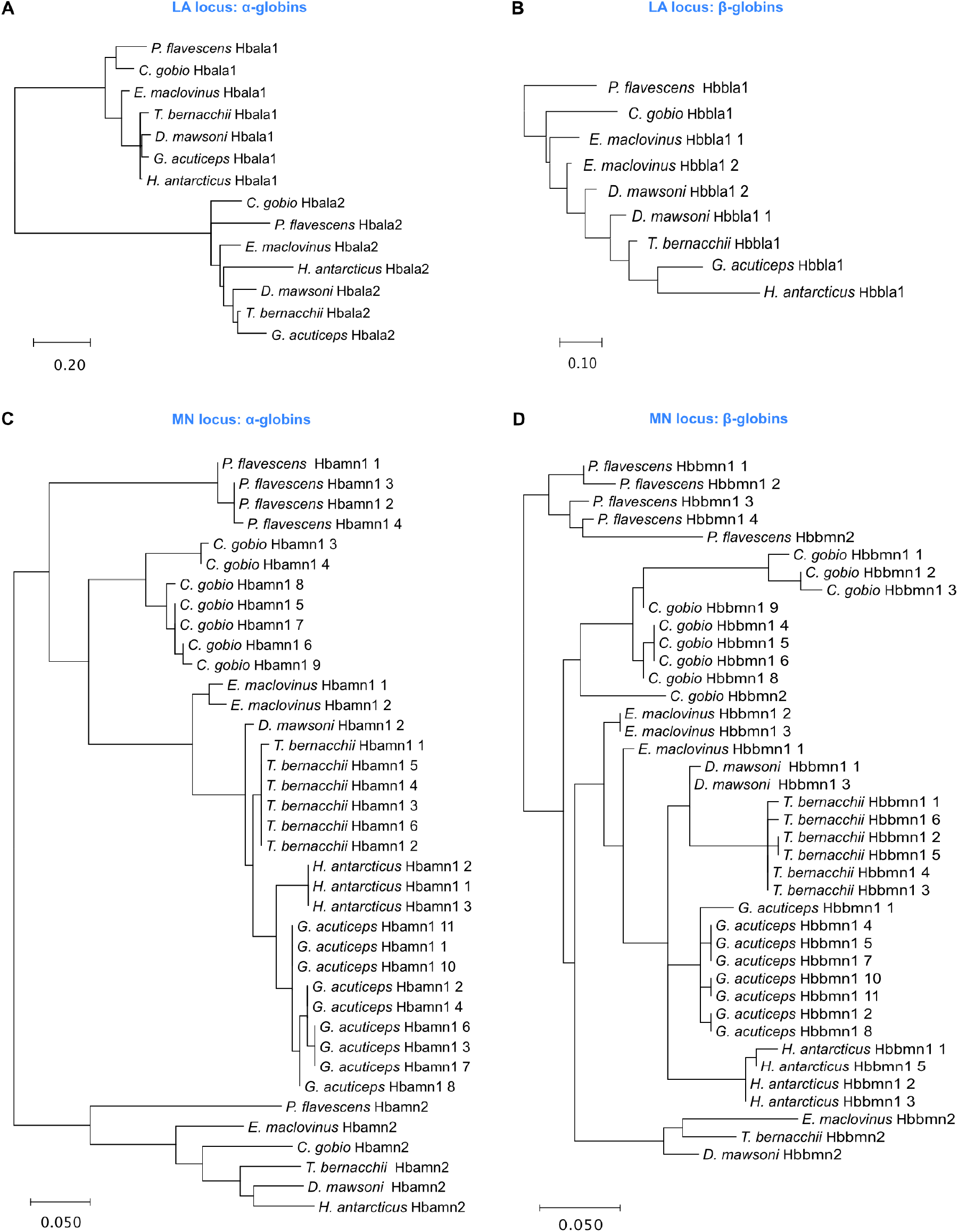
Haemoglobin protein trees (RaxML) from both clusters: A) α-globins in LA locus, B) β-globins in LA locus, C) α-globins in MN locus, D) β-globins in MN locus. The protein names are shown as Hbamn, Hbala for α-globins in MN and LA locus, and Hbbmn, Hbbla, for β-globins in MN and LA locus, followed by the number of the specific copy (for naming convention used see **Methods**, and **Supplementary Methods**).

**Extended Data Figure 7.**
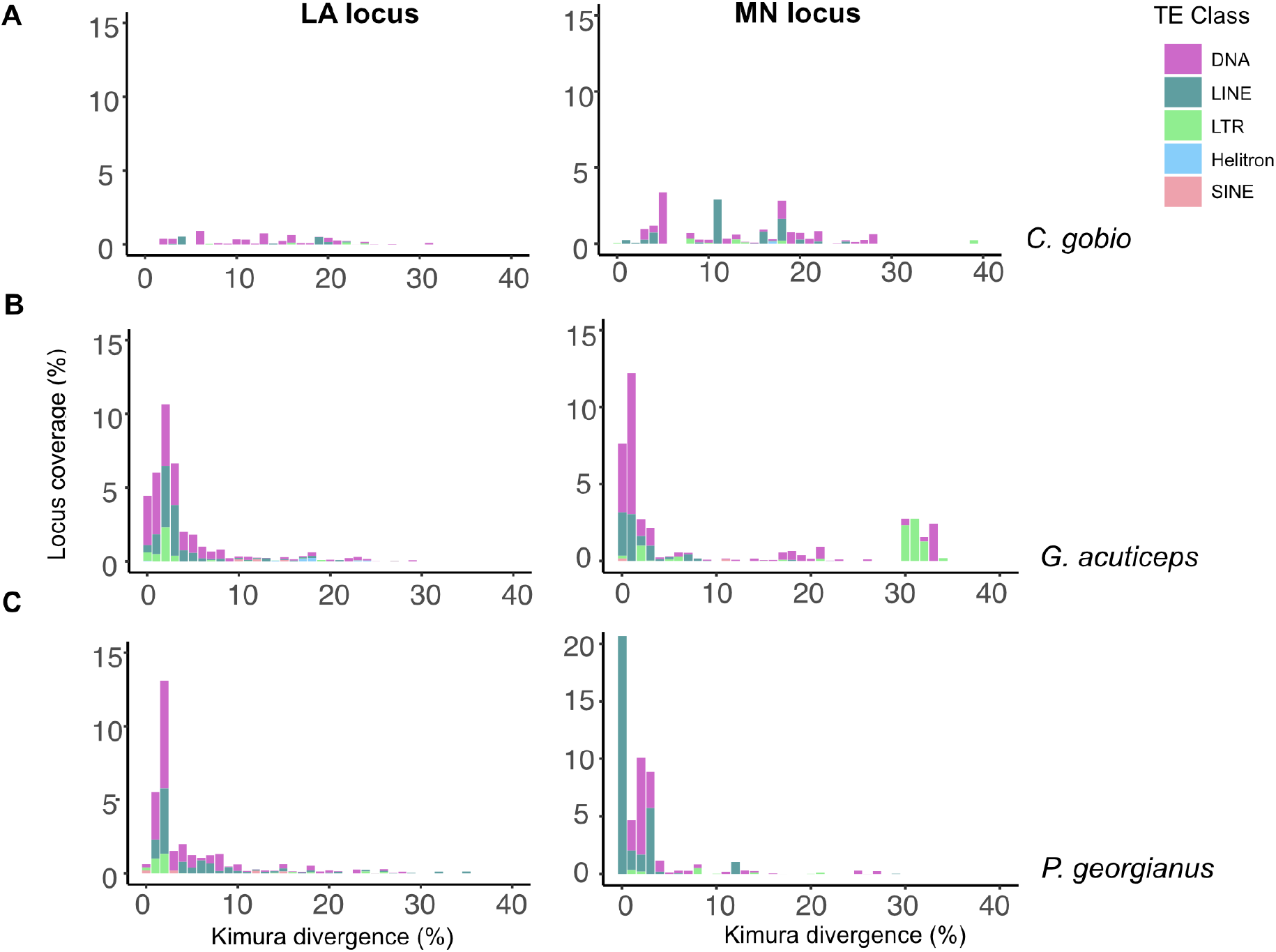
Repeat landscapes (as in **Extended Data Figure 5**) of transposon insertions in haemoglobin clusters LA and MN: A) *C. gobio*, B) *G. acuticeps*, and C) *P. georgianus*. The colours represent different TE classes.

## Acknowledgements

We thank the Wellcome Sanger Institute Scientific Operations for help with sequencing data production. We thank Laura Gerrish for help with a map of Antarctica and Southern Ocean. I.B., S.A.M, R.D. were supported by Wellcome grants WT207492 and WT206194; I.B. and E.A.M. by Wellcome grants 104640 and 092096; C.H.C.C. by US National Science Foundation grant ANT11-42158; M.M. by a mobility fellowship from the Norwegian Research Council (FRIPRO 275869); T.D., J.H.P by NSF OPP-1543383 and OPP-1947040; H.W.D. by US National Science Foundation grants OPP-0132032, PLR-1444167, and OPP-1955368; M.S.C. by NERC-UKRI core funding to the British Antarctic Survey; W.S. by Swiss National Science Foundation (176039). J.M.D.W., Y.S., J.T., W.C., K.H. by Wellcome WT206194; L.H. by WT108749 and WT222155.

IB dedicates this work to the memory of her father CB.

## Data Availability

The data generated in this project have been deposited on NCBI under BioProject PRJEB53202 and the following accessions: GCA_900634415, GCA_902827165, GCA_902827135, GCA_902827175, and GCA_902827115. For the purpose of open access, the author has applied a CC BY public copyright licence to any Author Accepted Manuscript version arising from this submission.

## Author contributions

I. B., R.D., E.A.M., H.W.D, J.H.P., C.H.C.C., M.S.C., T.D., W.S. designed the study and selected species for sequencing

I.B. led design, data generation, and analysis

H. W.D., C.H.C.C., J.H.P., T.D., M.S.C. contributed samples

I. B., M.S., K.O. generated sequencing data

I. B., S.A.M., Z.N. assembled genomes

J. M.D.W., A.T., Y.S., J.T., W.C., K.H. performed genome curation

I.B., T.D., M.M., C.H.C.C., J.M.D.W. performed data analyses

I.B. performed transposon annotation

L.H. performed gene annotation

I.B., J.T. prepared data for submission

R.D., E.A.M. provided computational resources and funding

I.B. wrote the manuscript, with edits from R.D., C.H.C.C., J.M.D.W., M.S.C., T.D., M.M., and comments from all authors

All authors reviewed the final manuscript and approved it.

