## Supplementary Text for "Genomics of cold adaptations in the Antarctic notothenioid fish radiation"

### Bista et al. Supplementary Text

#### Supplementary Results

##### Sequencing and assembly

We generated and analysed reference genome sequences for 24 species, across the entire notothenioid radiation, of which 11 species were assembled with 10X Genomics data and 8 species assembled using only Illumina HiSeqX reads.

We assembled the 10X Genomics sequenced species with Supernova 2.0[1], by sequencing a single linked-read library per species, with average 741 million reads per species, and average molecule length 45 kb (**Supplementary Table ST9**). Average continuity (N50) was 2,637,607 kb, ranging between 59,550 kb (*Histiodraco velifer*) and 14,0721,78 kb (*Bovichtus variegatus*), with an average 46,494 number of scaffolds (11,172 *Bovichtus variegatus* to 83,165 *Akarotaxis nudiceps*) (**Supplementary Table ST2**). For the 8 species with only Illumina HiSeq reads we generated as primary assembly with SOAPdenovo2[2] scaffolded using synteny with the most closely related PacBio-based genome assembly (with only exception *Bathydraco marri* for which scaffolding failed*).* This approach generally improved the N50 and BUSCO completeness for these species compared to the primary. BUSCO gene completeness averaged (95%) for PacBio assemblies, (86%) for Supernova assemblies, and (64%) for SOAPdenovo-based assemblies.

##### Phylogenetic analysis

The BEAST2 analysis of concatenated datasets presented in **Fig. 2A** was supported by the alternative phylogenetic approaches employing IQ-TREE and ASTRAL. The tree topologies inferred from the strict, permissive, and full sets of alignments were identical to the topology inferred with BEAST2, except for the positions of *G. acuticeps* and the outgroup species *Takifugu rubripes*. *G. acuticeps* was inferred to be the sister lineage to Channichthyidae in the IQ-TREE analyses of the concatenated permissive and full dataset, respectively; however, the species was grouped with the other representatives of Bathydraconidae in the IQ-TREE analysis of the strict dataset, just like in the BEAST2 analysis. The node grouping *G. acuticeps* with Channichthyidae in the two analyses was poorly supported, with bootstrap values of 82 and 70, and with site-concordance factors[3] of 18.2 and 18.0 in the analyses with the permissive and full datasets, respectively. The position of *T. rubripes* outside of Perciformes received bootstrap support values between 87 and 98, and site concordance factors between 16.4 and 22.9, in the IQ-TREE analyses with the three datasets. Besides these two problematic species, the rest of the topology was generally strongly supported, with mean bootstrap values between 97.0 and 98.6, and mean site-concordance factors between 57.5 and 57.9.

Analysing each alignment separately with IQ-TREE led to mean gene-concordance factors[3] between 50.8 and 54.5. The gene-concordance factors for the node combining *G. acuticeps* and Channichthyidae were 36.7 and 36.8 in the analyses with the permissive and full dataset, respectively. The node combining *G. acuticeps* with Bathydraconidae in the analyses with the strict dataset received comparable bootstrap, site-concordance, and gene-concordance values (53, 18.8, and 36.6).

When using the gene trees inferred with IQ-TREE as input for species-tree analyses with ASTRAL, the same species tree was inferred with all three datasets, and this tree was identical to the one inferred with IQ-TREE based on the permissive and full datasets. In these analyses, the position of *G. acuticeps* with Channichthyidae was supported by Bayesian posterior probabilities between 0.38 and 0.62. All other nodes received far greater support in the ASTRAL analyses, with mean Bayesian posterior probabilities between 0.97 and 0.98.

#### Supplementary Methods

##### Genome assembly

The assembly for ***Cottoperca gobio*** (channel bull blenny) is based on 75x PacBio Sequel data, and 54x Illumina HiSeqX data generated from a 10X Genomics Chromium library, as well as Bionano Saphyr two-enzyme data generated with Bionano Irys, generated at the Sanger Institute. Additionally, 145x coverage HiSeqX data from a Hi-C library prepared by Arima Genomics, using tissue from a different individual (fCotGob2) than the other data. An initial PacBio assembly was made using Falcon-unzip[4] without repeat-masking during overlap detection. The primary contigs were first scaffolded using a wtdbg assembly as a guide, then scaffolded further using the 10X data with scaff10x and then with Bionano two-enzyme hybrid scaffolding. After using the PacBio data to gap-fill with PBJelly and polish with Arrow, the assembly was polished again using the 10X Illumina data and freebayes. Contiguity was then further increased by filling gaps with the contigs from a wtdgb assembly made from Canu[5] corrected PacBio reads. The assembly was then re-polished with Arrow and freebayes, and retained haplotigs were identified with purge_haplotigs[6]. Finally, the assembly was scaffolded to chromosomes using Arima Hi-C data and manually improved using gEVAL[7] to correct mis-joins and improve concordance with the Bionano data and Arima Hi-C data. The chromosomes were named based on synteny to the Japanese medaka assembly (*Oryzias latipes*) GCA_002234675.1.

The assembly for ***Trematomus bernacchii*** (emerald notothen), was based on 46x PacBio data and 53x of 10X Genomics Chromium data. The assembly process included the following sequence of steps: initial PacBio assembly generation with Falcon-unzip, retained haplotig identification with purge_haplotigs, 10X based scaffolding with scaff10x, Arrow polishing, and two rounds of FreeBayes polishing. The assembly was analysed and manually improved using gEVAL. A second round of haplotig retention was run on the curated assembly using purge_dups [8].

The assembly for ***Harpagifer antarcticus*** (Antarctic spiny plunderfish), was based on 67X PacBio data, 40x of 10X Genomics Chromium data, and Bionano Irys data. The assembly process included the following sequence of steps: initial PacBio assembly generation with Falcon-unzip, retained haplotig identification with purge_haplotigs, 10X based scaffolding with scaff10x, Bionano hybrid-scaffolding, Arrow polishing, and two rounds of FreeBayes polishing. The assembly was analysed and manually improved using gEVAL. A second round of haplotig retention was run on the curated assembly using purge_dups [8].

The assembly for ***Gymnodraco acuticeps*** (Ploughfish), was based on 31x PacBio data, and 41.8x of 10X Genomics Chromium data. The assembly process included the following sequence of steps: initial PacBio assembly generation with Falcon-unzip, retained haplotig identification with purge_haplotigs, 10X based scaffolding with scaff10x, Arrow polishing, and two rounds of FreeBayes polishing. The assembly was analysed and manually improved using gEVAL. A second round of haplotig retention was run on the curated assembly using purge_dups [8].

The assembly for ***Pseudochaenichthys georgianus*** (South Georgia icefish), was based on 92.9x PacBio data, 56x 10X Genomics Chromium data, and Dovetail Hi-C data generated at the Wellcome Sanger Institute. The assembly process included the following sequence of steps: initial PacBio assembly generation with Falcon-unzip, retained haplotig identification with purge_haplotigs, 10X based scaffolding with scaff10x, Bionano hybrid-scaffolding, Hi-C based scaffolding with SALSA2, Arrow polishing, and two rounds of FreeBayes polishing. The assembly was analysed and manually improved using gEVAL. A second round of haplotig retention was run on the curated assembly using purge_dups [8].

#### Assembly curation

Manual curation can substantially improve the continuity and accuracy of genome assemblies[9]. In order to further improve the quality of these genomes we performed manual curation to remove miss-assemblies, duplications, sequencing contamination and introduce joints based on supporting evidence. Each assembly was manually curated[10] using the Genome Evaluation Browser (gEVAL)[7]. Initially contig and scaffold integrity was confirmed with PacBio read mapping and 10X illumina read mapping. Scaffolding was enhanced using 10X read information and contig end sequence overlaps. In addition, for fHarAnt1 scaffold integrity was further confirmed using Bionano BssSI optical maps, visualised in Bionano Access, breaking and re-joining where necessary. For fPseGeo1 a 2D map was built using Hi-C reads and visualised on HiGlass[11], which allowed for the further scaffold correction and super scaffolding in order to bring the assembly to chromosome scale. Chromosome name assignment was based on comparative alignment to the fCotGob3.1 chromosome assembly GCA_900634415.1. The curated assemblies were finally run through purge_dups[8] to remove artificially retained haplotypic duplication (**Supplementary Table ST10**).

#### RNAseq

Transcriptome data were generated to improve gene annotation. Total RNA for RNAseq was extracted using the RNeasy Qiagen extraction kit, from approximately 20-40mg of tissue. The RNA quality was assessed with the Qubit HS RNA kit and Agilent Bioanalyzer Nano chips, and only extracts with RIN value >8 were used for sequencing. Sequencing was performed on Illumina HiSeqX, for a variety of tissues. For *C. gobio* four tissues used included brain, muscle, and gonads from one individual, preserved in RNAlater, and frozen spleen from a different individual. For *T. bernacchii* four tissue types were used for RNAseq including brain, muscle, ovary, and testis, preserved in RNAlater. For *G. acuticeps* two tissues were used for RNAseq, including brain and ovary from individual, preserved in RNAlater.

#### Gene annotation

The gene sets for *C. gobio, T. bernacchii*, *H. antarcticus, G. acuticeps,* and *P. georgianus* were generated via the Ensembl Gene Annotation system[12], and are available as part of Ensembl and Ensembl Rapid release (rapid.ensembl.org). Annotation was created primarily through alignment of short read RNA-seq data to the genome. The RNA-seq short-read data were sourced from samples generated as part of projects with the following BioProject IDs: PRJEB26835, PRJNA263718, PRJNA287661, PRJNA308624, PRJNA401363, PRJNA471228, PRJEB25429, PRJNA301149 and PRJNA422913. Gaps in the annotation were filled via protein-to-genome alignments of a select set of vertebrate proteins from UniProt[13], which had experimental evidence for existence at the protein or transcript level.

At each locus, the data were collapsed and consolidated, with priority given to models derived from the RNA-seq data, producing a set of final gene models along with their associated non-redundant transcript set. To help differentiate between true isoforms and fragments, the likelihood of each ORF was assessed in relation to known vertebrate proteins. Low quality transcript models, e.g. those with evidence of a fragmented ORF, were removed. In loci where the RNA-seq data were fragmented or missing, homology data took precedence, with preference given to longer transcripts that had strong intron support from the short-read data.

Gene models from the above process were classified into three main types: protein-coding, pseudogene, and long non-coding. Models with hits to know proteins, and few structural abnormalities were classified as protein-coding. Models with hits to known proteins that also display abnormalities such as absence of a start codon, non-canonical splicing, unusually small intron structures (< 75bp) or excessive repeat coverage, were reclassified as pseudogenes. Single-exon models with a corresponding multi-exon copy elsewhere in the genome were classified as processed (retrotransposed) pseudogenes. If a model failed to meet the criteria of any of the previously described categories, did not overlap a protein-coding gene, and had been constructed from transcriptomic data then it was considered as a potential lncRNA. Potential lncRNAs were additionally filtered to remove single-exon loci due to the unreliability of such models.

Putative miRNAs were predicted via a BLAST of miRBase[14] against the genome, before passing the results to RNAfold[15]. Other small non-coding loci were identified by scanning Rfam[16] against the genome (described in more detail in[12] ) and passing the results into Infernal[17].

#### New naming system for teleost haemoglobin genes

The current haemoglobin gene naming system in fish mostly relies on the zebrafish laboratory model species and on the expression pattern of each of its haemoglobin genes during embryonic and/or adult phases. While informative for zebrafish research, using a naming system based on embryonic or adult expression for species in which expression dynamics of haemoglobin genes can't be assessed may lead to misinterpretations, especially because expression patterns of haemoglobin genes are known to be influenced by local organisation of the genomic region that may not be conserved across species[18,19]. Therefore, designating orthologous haemoglobin genes across species needs a nomenclature system that is independent of an expression pattern that may not be evolutionarily conserved. We thus propose here a novel naming system based on genomic organisation rather than expression data.

First, the established haemoglobin alpha and beta denominations (i.e., *hba* and *hbb*) are conserved due to clear sequence conservation. Second, the presence of each gene in the LA or the MN cluster is added as a suffix (e.g., *hbala* and *hbbmn*). Third, a final numeral suffix is added to reflect the relative positioning of the gene within each cluster. The orientation of the most upstream *hba* gene determines the orientation of the cluster and is arbitrarily named *hbala1* and *hbamn1* for the LA and MN clusters, respectively. Thus, the nomenclature reflects position but not necessarily orthology. The neighbouring *hbb* gene is called *hbbla1* and *hbbmn1* for the LA and MN clusters, respectively. The names of genes further to the conventional right of the locus are suffixed with incremental numbers following the orientation of the cluster. Tandem duplicated genes (e.g., *hbamn1.1* and *hbamn1.2*) and pseudogene (e.g., *hbamn1.3p*) nomenclatures follow zebrafish gene nomenclature guidelines https://zfin.atlassian.net/wiki/spaces/general/pages/1818394635/ZFIN+Zebrafish+Nomenclature+Conventions) established by ZFIN[20].

#### Phylogenetic analysis

Amino-acid sequences for 266 selected BUSCO genes were aligned with MAFFT v.7.453[21]. The 266 alignments were inspected by eye, and apparently misaligned sequence regions were set to missing data. A total of 1,141,524 amino acids were set to missing out of 6,410,688, including nine alignments that were excluded completely, leaving 257 alignments for further analysis. We then aligned nucleotide sequences of the same BUSCO genes according to the amino-acid alignments, ensuring that regions corresponding to the removed sequences were again set to missing data in the nucleotide sequence alignments. Sites with high entropy (entropy like score > 0.5) or high proportion of missing data (gap rate > 0.2) were removed with BMGE v.1.1[22], and alignments with more than three completely missing sequences, a minimum length below 500 bp, or a standard deviation of among-sequence GC-content variation greater than 0.03 were excluded. These filters were passed by 228 alignments.

Per alignment, we performed gene-tree analyses with the program BEAST2 v.2.6.0[23], with a Markov-chain Monte Carlo chain length of 25 million iterations, assuming the Yule model of diversification[24] and the uncorrelated lognormal relaxed clock model[25], and averaging over substitution models with the bModelTest add-on package[26]. These gene trees were time-calibrated by arbitrarily constraining their root age to 100 million years (with a standard deviation of 0.1). Chain convergence was suggested by effective sample sizes (ESS) per parameter greater than 200.

Based on the minimum ESS value per alignment and estimates for the mutation rate and its among-species variation, we identified the most suitable alignments for further phylogenomic analyses. We compiled a "strict" set of alignments that included all those that had a mean mutation rate estimate below 0.002 per bp per million year, a mutation rate standard deviation (relative to the mean estimate) below 0.9, and a minimum ESS value greater than 100; this set was a subset of a second, "permissive" set of alignments in which we placed those that had a mean mutation rate estimate below 0.00025 per bp per million year, a mutation rate standard deviation below 1.1, and a minimum ESS value greater than 50. The strict and permissive sets contained 140 and 200 alignments, respectively.

For the strict set of 140 alignments, the permissive set of 200 alignments, and the "full" set of 257 alignments, we performed maximum-likelihood phylogenetic analyses with IQ-TREE v.1.7[27] after alignment concatenation, maintaining separate partitions with unlinked instances of the GTR+Gamma substitution model for each of the original alignments. Node support was assessed with 1,000 ultrafast bootstrap replicates[28]. Each of the three analyses was complemented with an estimation of gene- and site-specific concordance factors[3], and the three resulting sets of gene trees were used for separate species-tree analyses with ASTRAL v.5.7.3[29].

Finally, we estimated the phylogeny and the divergence times of notothenioid species with BEAST2 from a concatenated alignment combining all alignments of the strict set. To avoid potentially saturated sites, we excluded all third codon positions from this analysis, and to reduce its computational demand we grouped 280 original data blocks (separating first and second codon positions for each of the 140 original alignments of the strict set) into 12 positions, selected with the rcluster algorithm of PartitionFinder v.2.1.1[30], assuming linked branch lengths, equal weights for all model parameters, a minimum partition size of 5,000 bp, and the GTR+Gamma substitution model. The same substitution model was also assumed in the BEAST2 analysis, together with the birth-death model of diversification[31] and the uncorrelated lognormal relaxed clock model[25]. Time calibration of the phylogeny was based on four age constraints defined according to a recent timeline of teleost evolution inferred from genome and fossil information[32]. The age of the most recent common ancestor of Eupercaria was constrained to around 97.47 MYA (2.5-97.5 interpercentile range: 91.3-104.0 Ma), that of the clade combining Eupercaria, Ovalentaria, and Anabantaria (without Carangaria as no members of Carangaria were included in our dataset) was constrained to around 101.79 Ma (95.4-109.0 MYA), that of the clade combining these four groups with Syngnatharia and Pelagiaria was constrained to around 104.48 Ma (97.3-112.0 Ma), and that of the clade combining those six groups with Gobiaria was constrained to around 107.08 (100.0-114.0 MYA). All constraints were implemented as lognormal prior distributions with mean values as specified above and a standard deviation between 0.033 and 0.036. Additionally, we constrained the monophyly of the groups Notothenioidei, Perciformes, Ovalentaria, Anabantaria, and the clade combining the latter two groups. All of these constraints agreed with findings of recent genomic studies of teleost relationships[32–35]. We performed six replicate BEAST2 analyses with lengths of 330 million MCMC iterations, of which the first 10% were considered as burn-in. Convergence among MCMC chains was confirmed by ESS values greater than 120 for all model parameters and greater than 270 for the likelihood and the prior and posterior probabilities. The posterior tree distribution was summarised in the form of a maximum-clade credibility tree with TreeAnnotator v.2.6.0[36]. We attempted to repeat the BEAST2 analyses with the permissive and full datasets, however, these analyses were too computationally demanding, so that even after 330 million MCMC iterations and run times of several months, some of the ESS values remained below 100. Nevertheless, the preliminary results from these analyses supported the same tree topology as the analyses with the strict dataset.
